# A Closed-Loop Robot Scientist for Autonomous Biological Discovery

**DOI:** 10.64898/2026.09.11.751076

**Authors:** Kameron Bielawski, Krishna Srinivasan, Nate Gaylinn, Shawn Beaulieu, Robert Brucker, Michael Levin, Jeantine Lunshof, Joshua Bongard, Douglas Blackiston

**Affiliations:** Department of Computer Science, University of Vermont, Burlington VT, USA; Department of Biology, Tufts University, Medford MA, USA; Wyss Institute for Biologically Inspired Engineering, Harvard University, Boston MA, USA; Department of Genetics, Harvard Medical School, Boston MA, USA; Vermont Complex Systems Institute, Burlington VT, USA

**Keywords:** Artificial intelligence, forward model, automation, organoid, embryo

## Abstract

Biological systems respond to a wide range of physical inputs and an overarching goal of biological research is understanding how these signaling events coordinate cellular, tissue, and organism-level outcomes. Yet the vast combinatorial space of physical and chemical interventions that influence these processes remains largely unexplored. Thus, a search process is required that efficiently learns how inputs affect a biological target via a series of automatically generated interventions, and a robot scientist that can conduct them. To this end we here introduce the Multimodal Organismal Modulation Robot (MOMbot), a robot scientist that integrates four physical intervention modalities—chemical delivery, electrical field application, mechanical vibration, and thermal modulation—within a single hardware platform. MOMbot’s multimodal hardware is paired with high-resolution imaging and an online active learning algorithm that autonomously generates, executes, and refines interventions based on accumulated data. We validate the system across biological materials spanning three orders of magnitude in scale, including *Xenopus laevis* embryos, motile mucociliary organoids, and disaggregated ectodermal stem cells, to show the utility of this approach across diverse biological disciplines. Using these biological models, we demonstrate precise thermal control of embryonic developmental rate, automated chemical perturbation with behavioral readouts, programmable electrical field stimulation, and vibrational assembly of functional mucociliary organoids from loose cells. We also describe the active learning algorithm that autonomously concentrates interventions near regions of maximal outcome uncertainty, yielding sample-efficient discovery of an electrical duration–response relationship for motile organoids. These results establish MOMbot as a scalable robot/AI scientist team for biological discovery, enabling autonomous, multimodal physical experimentation and AI-driven exploration of complex biological systems.

## 1 Introduction and Background

Disembodied AI—computational models and data analysis algorithms operating on existing datasets—has already transformed biological research [1]. However, biological discovery ultimately requires physical interaction with living matter, and over the past two decades, embodied AI in the form of intelligent robotic experimentation systems has become a prominent collaborator in laboratories globally. In all basic sciences, closed-loop experimentation has been achieved by coupling automated experimental design and learning algorithms to physical hardware to execute experiments. In chemistry [2–6], physics [7, 8], materials science [9–13], molecular biology [14, 15], and many other subdisciplines [16–18], scientists and engineers have implemented scientifically fruitful autonomous experimentation systems. Most relevantly, in biology, these systems have been used to optimize cell culture media composition [19], optimize cell culture protocols [20], evaluate the cognitive capabilities of model organisms [21, 22], perform tissue scratch assays and evaluate collective migration [23], and design polymer-protein hybrids [24]. While these approaches have been proven successful among specific fields of experimentation, there remains a lack of turnkey systems which would enable discovery across several scales of biological organization.

Physical interventions can, in principle, also deflect biological system dynamics toward new, stable phenotypes more easily than top-down design at the genetic or molecular level. But closing this gap with other branches of science already automated requires systematic, high-throughput and efficient exploration of the response landscape of diverse biomaterials, accelerating fundamental understanding of biological development while simultaneously providing a programmable interface for synthetic biology and bioengineering— the directed construction of engineered living systems through real-time physical modulation of cellular self-organization.

To address this, we sought to close the automatic experimentation loop for physical modulation of biological systems by building a system that can both deliver diverse physical interventions and autonomously decide which interventions to try next. Here we present a Multimodal Organismal Modulation Robot (MOMbot), a new integrated hardware system controlled by an online artificial intelligence platform to autonomously choose optimal experiments given human-specified research or bioengineering objectives. MOMbot is a scalable laboratory robot which can programmatically deliver four types of physical stimulus (thermal, electrical, mechanical, and chemical) simultaneously to biological material within five independent experimental stations, each with two dishes, where each dish is equipped with a high-resolution imaging system to observe experimental results. Practically, this system can scale up the rate of information-dense experimentation by both increasing throughput with multiplexed experiment stations and by using AI to automatically discover and execute experiments with high expected information content based on past observations.

The combination of high resolution optics and large culture environment enables experimentation on biomaterials spanning several orders of magnitude. We demonstrate this by conducting experiments on developing *Xenopus* embryos, motile mucociliary organoids, and disaggregated ectodermal stem cells. These varied materials and spatial scales highlight the broad potential of this system across multiple areas of biomedical and basic sciences. Vertebrate embryos continue to be foundational models for uncovering the mechanisms underlying morphogenesis, developmental defects, and stem cell differentiation [25–27]. In parallel, amphibian mucociliary organoids and cell based approaches have proven to be a powerful model for understanding human airway diseases [28–30], as the cell types, emergent self-organization, and tissue level function is highly conserved between the two systems. Beyond these biomedical contexts, materials science and biorobotics approaches leverage the native properties of living cells toward novel bioengineering goals, like the harnessing of self-organizing dynamics [31, 32], functional synthetic-biotic integration [33–35], and the construction of novel morphologies [36–38].

A challenge in biological research remains the hyperspace of experimental variables that can be investigated in tandem. Beyond transcriptional networks which can be interrogated with genetic and pharmacological approaches, many cells and tissues also respond to mechanical cues, vibroacoustic stimuli, temperature, exogenous or endogenous electric fields, pH, reactive oxygen species, and other physiological signals, to name only a subset. Tuning these stimuli in real-time, alone or in combination, incurs significant labor cost, and existing automated platforms implement only a subset of these stimuli—often in experimental environments not suitable for both cells and organism-scale experiments, as in the case of microfluidic systems [39, 40]. Moreover, the majority of automation requires manual stimulus scheduling by a human operator [41, 42], limiting throughput even when hardware is available. Closed-loop machine learning approaches can address both challenges: not only automating stimulus delivery, but also meaningfully participating in experimental design through Bayesian optimization and active learning techniques [43–47]. Rather than simply fitting models to existing data, active learning algorithms strategically select the most informative experiments to perform. By prioritizing information-dense samples, these methods can achieve strong model performance with relatively small datasets—an important advantage in laboratory settings where data collection is costly and experiments are time-intensive [48]. The active learner we deploy here has a minimal proof-of-principle design, which seeks to execute experiments whose outcomes are highly uncertain, and thus sample efficiently elucidates a duration-response curve.

In the proceeding, we instantiate this framework in hardware and software. We first present the physical architecture of MOMbot, detailing its multimodal stimulus integration, imaging infrastructure, and data management pipeline. We then report four experiments that collectively validate each intervention modality—vibrational assembly of functional organoids from dissociated cells, automated chemical perturbation of motile organoids, thermal modulation of embryonic developmental rate, and a proof-of-principle closed-loop active learning campaign targeting an electrical to cellular duration–response relationship. Together, these demonstrations establish both the physical versatility of the platform and the feasibility of autonomous, information-seeking experimentation in biological discovery.

## 2 Results

### 2.1 System Workflow

The following is a description of MOMbot’s experimental workflow from the perspective of a wet-lab scientist (herein the MOMbot “operator”). After sample preparation (see Methods 4.3), a human operator loads biomaterial into individual MOMbot dishes, which can be experimented upon individually or in parallel (Figure 1a). The operator then provisions the newly loaded dishes for the onboard software via a networked user interface, accessible locally or remotely, where the choice is made to expose the dish for AI-designed or human specified interventions. Following software initiation, images are collected at AI or human operator defined capture rates, and interventions are delivered via the station’s stimulus hardware with open or closedloop scheduling (Figure 1b). In the case of closed-loop scheduling, the remote AI observes the biomaterial in real-time, and uses observable behaviors to decide upon interventions (Figure 1c). Experimental data accumulates (Figure 1d) in both human operator and AI designed experiments, and upon completion of each experiment dishes are collected for disposal and replaced allowing the experimental pipeline to repeat as necessary.

**Figure 1:**
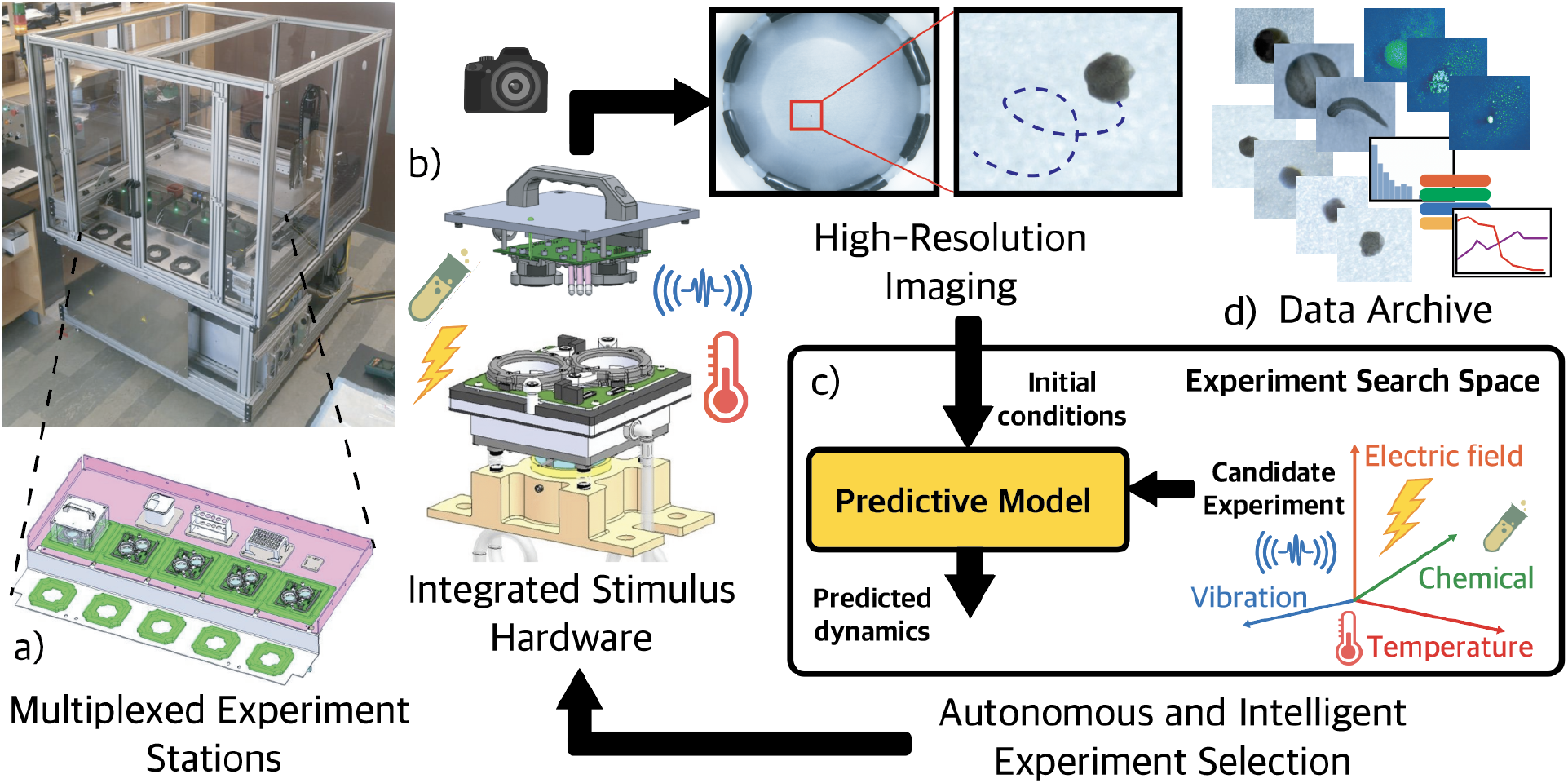
Architecture of the Multimodal Organismal Modulation robot (”MOMbot”). a) MOMbot is a laboratory hardware system with five experiment stations, each with two experiment dishes. The schematic view shows the station work zone (green) and chemical library/waste area (pink). A liquid handling system can access the entire experimental space, dispensing microliter volumes of chemicals to each dish as necessary.b) Each station integrates four stimulus modalities (chemical, vibration, electric fields, and temperature), as well as two high-resolution cameras for each pair of experiment dishes. c) The hardware is controllable by intelligent software active learning algorithms, enabling full autonomy in the selection and execution of experimental stimuli. d) A data archive accumulates dish-annotated data from all experiments executed on MOMbot.

### 2.2 Experimental Overview

Biological systems operate within a high-dimensional landscape of interacting stimuli, presenting a significant challenge for systematic experimental exploration. Towards overcoming this challenge, we demonstrate the hardware, software, and wetware capabilities of the MOMbot platform with four classes of experiments validating the integrated system architecture and active learner capabilities for autonomous experimentation. First, the vibroacoustic capabilities (Figure 2c) of the platform were validated by automatically and mechanically aggregating populations of loose *Xenopus laevis* ectodermal stem cells into dense clusters, which then form spheroids and self-organize into functional mucociliary organoids, demonstrating a novel autonomous organoid manufacturing method from uniformly distributed fields of cells (Figure 3). In the same experiment the single-cell resolution of the system’s imaging hardware (Figure 2a) is highlighted. Second, the system’s chemical delivery mechanism (Figure 2b) semi-automatically delivers a reactive oxygen species, H_2_O_2_, to motile organoids, whose behavioral response varies depending on dosage. Third, *Xenopus* embryonic development rate is modulated using the system’s temperature modulation (Figure 2e, Figure 4b). Fourth and finally, an active learning algorithm automatically carries out experiments on single self-motile organoids in a “hello, world” research agenda to discover the relationship between electrical stimulus and organoid velocity (Figure 1c). As a proof-of-principle, the active learning algorithm autonomously concentrates experiments near the region of maximal outcome uncertainty in the duration-response landscape, achieving more informative coverage of the response function than undirected uniform sampling of the parameter space (Figure 5).

**Figure 2:**
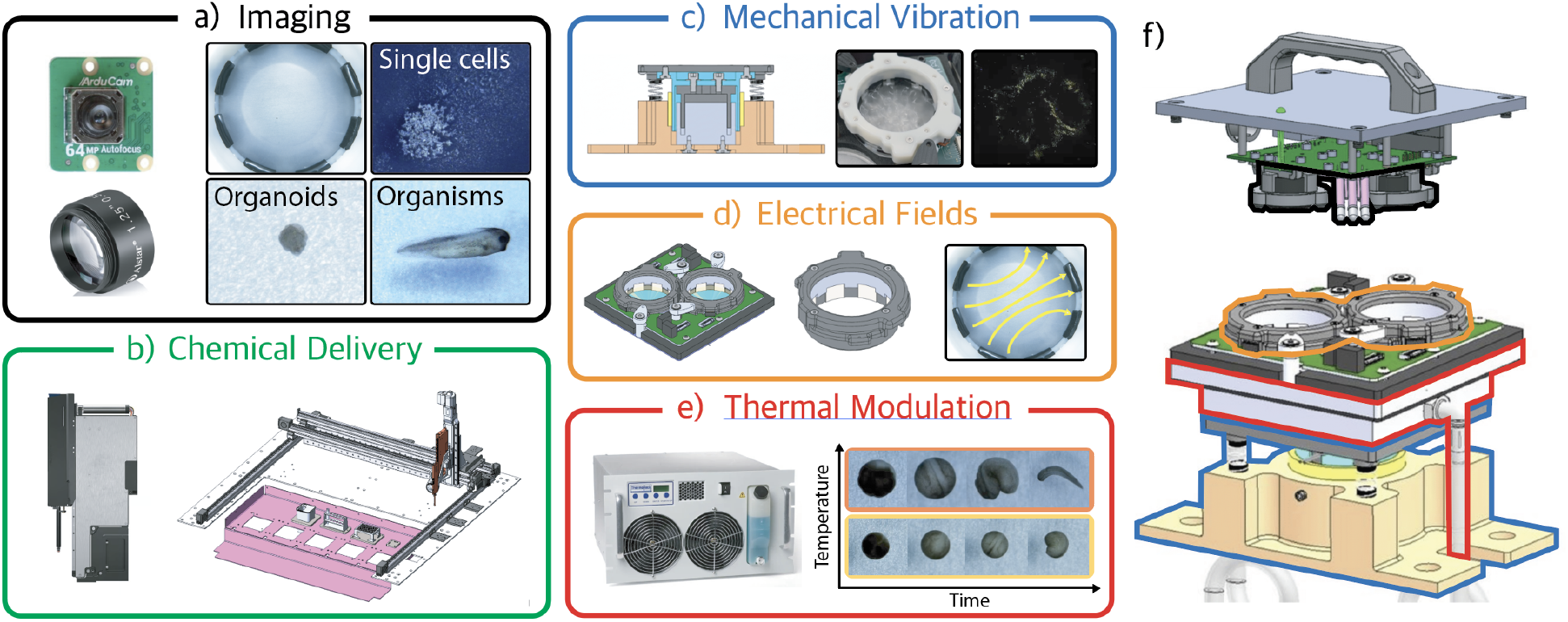
Experiment station and stimulus overview. a) Two high-resolution cameras sit above the centers of each experiment dish, enabling imaging of single cells, organoids, and mm to cm sized organisms (see Methods Section 4.1.2). b) MOMbot is equipped with a gantry carrying a liquid handling system, a pipette tip dispenser, chemical reservoirs, and a pipette tip discard bin, for semi-autonomous chemical delivery (see Methods Section 4.1.3). c) Each station is capable of mechanical vibrations in a range from 7-200Hz at user specified amplitudes (see Methods Section 4.1.4) (rightmost dark panel shows spatial patterning of cells due to vibration in MOMbot, similar to Chladni patterns). d) Dishes in each station support electrodelined inserts to independently modulate electrical fields of user defined cardinal directions and intensities (see Methods Section 4.1.6). e) Each station’s temperature can be independently modulated from 4C to 30C, enabling speeding and slowing of biological development (see Methods Section 4.1.5). f) All of these capabilities are integrated into a single hardware device, each with two experimental dishes (MOMbot houses 5 of these stations).

**Figure 3:**
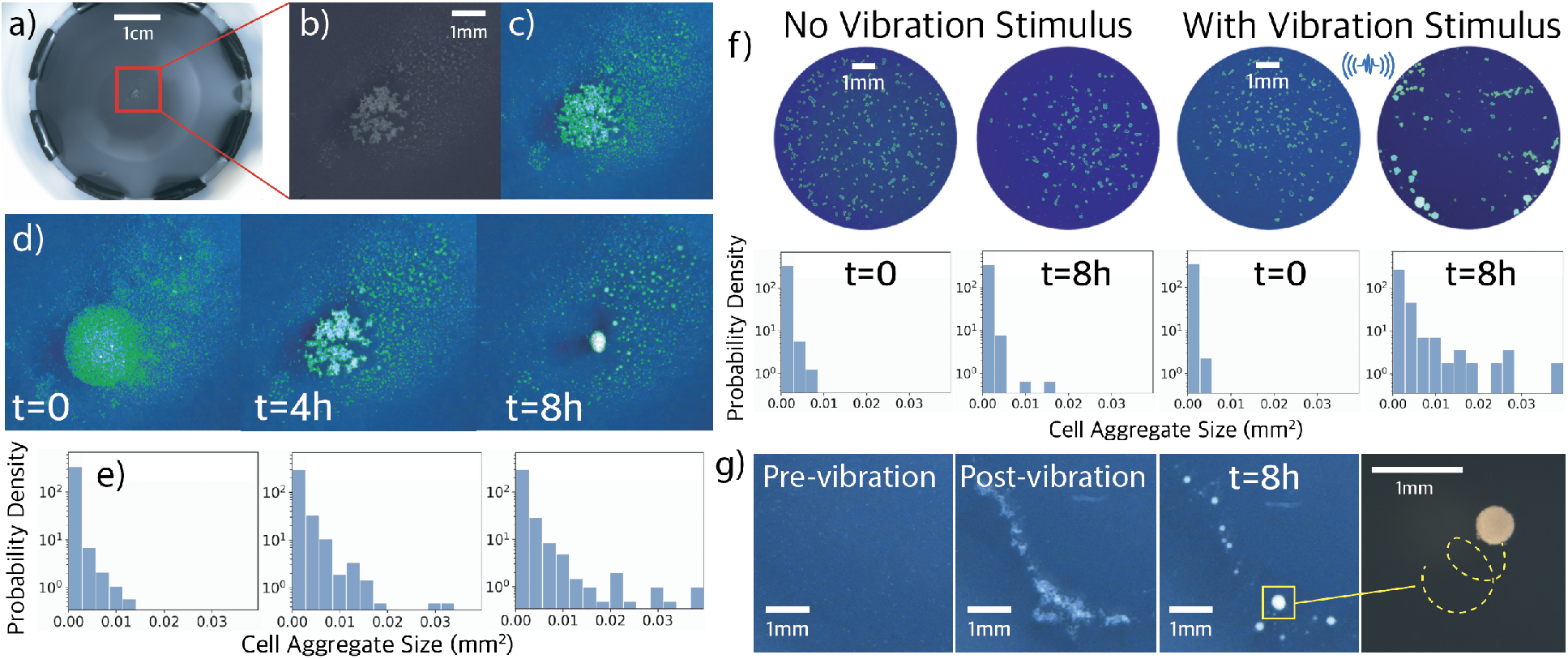
Imaging and manipulation of dispersed stem cell populations with vibroacoustic stimulation. a) Raw image from MOMbot dish. b) Cropped raw image of loose deep ectodermal *Xenopus* cells. c) Contrast-adjusted and segmented image, displaying single-cell segmentation ability. d) Cell aggregation over 8 hours. e) Distribution of cell aggregate sizes over time. f) Vibration validation study: starting with initially uniformly scattered loose cells (t=0), do nothing (middle-left, t=8h) as a control and repeatedly vibrate the cells into larger clusters where they can aggregate (right, t=8h). Observe differing distributions of aggregate sizes. g) Time sequence of vibration (left to right): pre-vibration uniformly distributed loose cells; post-vibration densely clustered cells; organoids form after aggregation; organoids become motile after many hours.

**Figure 4:**
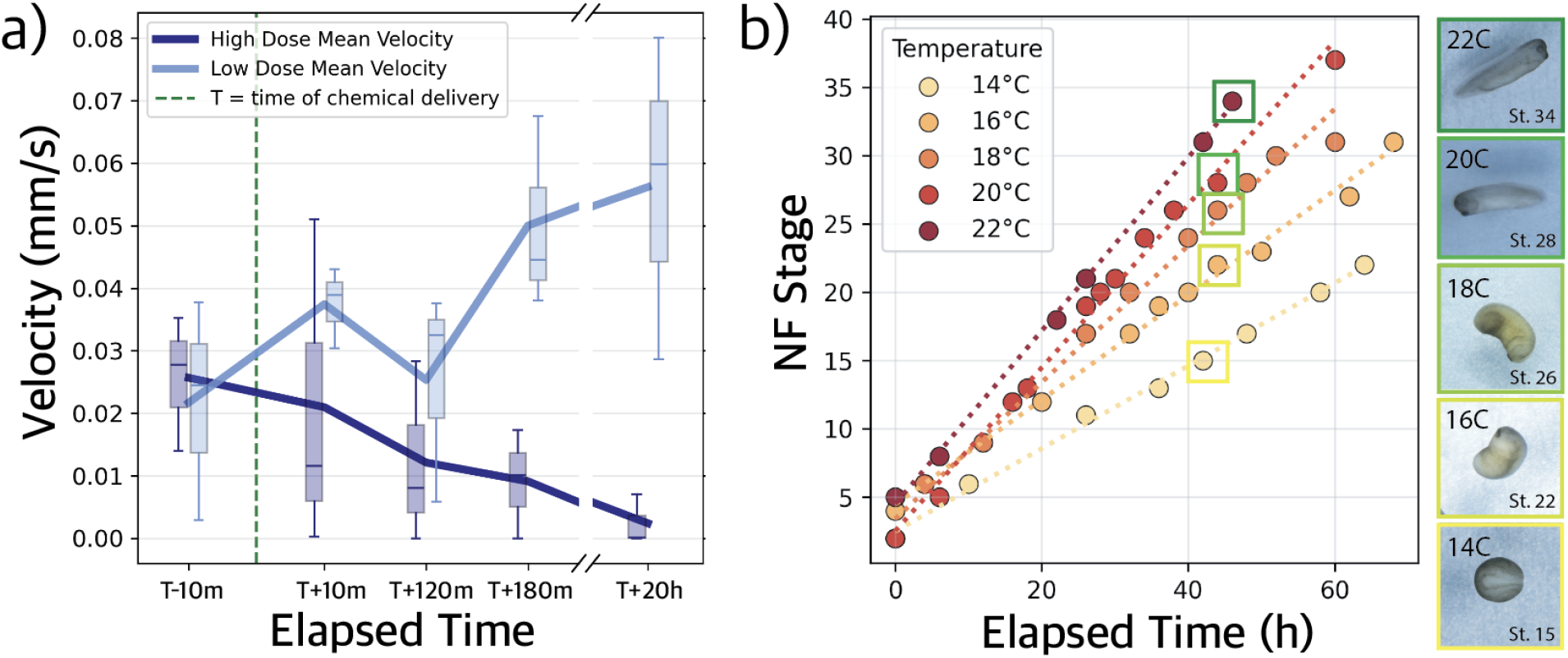
Control of tissue function and organism development with chemical and temperature stimulation. a) Individual mucociliary organoids were exposed to 98µM and 980µM H_2_O_2_. The higher concentration resulted in cessation of organoid movement, which is a proxy for motile ciliary beating. All organoids survived the duration of the experiment, changes in velocity were not a result of tissue death. b) Individual fertilized *Xenopus* embryos were raised under several temperatures known to be tolerated across embryogenesis. Individuals at 22°C developed at three times the rate of 14°C siblings, matching established baselines in the field. Dotted line indicates the time of H_2_O_2_ delivery. NF = Nieuwkoop and Faber stages.

**Figure 5:**
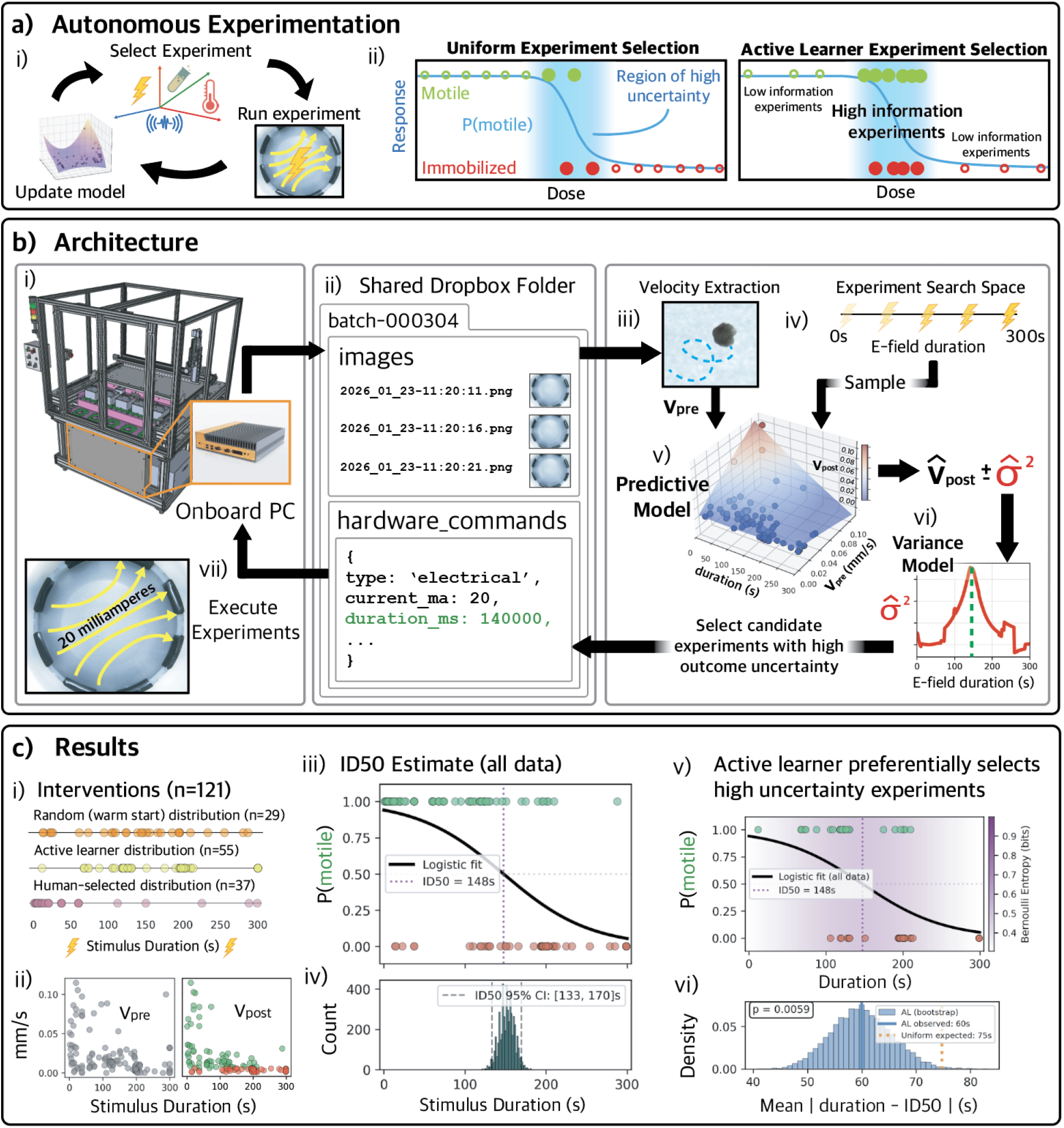
Overview of autonomous experimentation. a) i) Closed-loop experimentation involves iterative intervention selection, execution, and model updates. ii) An active learning based algorithm can seek out interventions with high expected information gain. b) Overview of experiment information flow. i) New biological material is loaded into a dish by an operator. ii) Images of the biological material are uploaded to a shared Dropbox folder which facilitates communication to and from MOMbot. iii) An image processing pipeline extracts an initial velocity from the raw data, and iv) the search space consisting of electric field durations is sampled. v) Candidate interventions and initial velocities are fed into a simple regression model to predict the outcome (post-intervention velocity) for a given initial condition/candidate intervention pair. vi) A variance model estimates the uncertainty of the predictive model’s estimate, and this variance informs the acquisition function which ultimately decides the candidate intervention chosen for execution on MOMbot. c) i) The total dataset accumulated on MOMbot consisted of *n* = 121 independent interventions conducted on motile organoids, where *n* = 29 electrical durations were randomly generated, *n* = 55 were generated by our algorithm, and *n* = 37 were generated by humans during system testing. ii) Organoid velocities prior to intervention (left), and after intervention (right); post-intervention immobilization was hand-labeled (right, red) if velocity=0 for the remainder of the observation period after the stimulus ended. iii) The “ID50” (Immobilization Duration 50)—the electrical stimulus duration at which 50% of organoids are immobilized—estimate generated from the entire dataset. iv) A bootstrap confidence interval estimate generated from uniform sampling validates the ID50 estimate. v) The *n* = 55 active learner’s samples were located nearer to the high uncertainty range than random sampling. vi) The average distance to the estimated ID50 of active learner samples (blue) is less than uniformly selected samples (orange) (*p <* 0.01, 1-sample t-test).

### 2.3 Hardware and Stimulus Validation

To validate the imaging capabilities of the system, populations of 15,000-20,000 deep ectodermal cells, collected and chemically disaggregated from developing *Xenopus laevis* embryos at the blastula stage (NF stage 9), were deposited into a dish containing 10ml of 0.75x Marc’s Modified Ringers (MMR). The system’s onboard optics can resolve single cells (Figure 3a–c), enabling segmentation and tracking of long-term aggregation dynamics. Across 8 successive hours of imaging, cells in contact with neighbors formed into aggregates of increasing size (Figure 3d–e), eventually forming three dimensional spheroids.

Vibroacoustic signals have been shown to have cell intrinsic effects via mechanosensitive and cytoskeletal mechanisms, as well as spatial applications via acoustic wells [49, 50]. To demonstrate the vibrational capabilities at the cell scale, dishes with loose embryonic ectodermal cells were loaded in a dispersed, uniformly distributed arrangement. One set of cells served as an untreated control, while the other received several iterations of vibrational stimulus at various frequencies between 40 and 70Hz via the onboard voice coil linear actuator (see Supplemental Materials for vibration parameters). Across the 8 hour observation period, untreated cells remained scattered in a dispersed field, while cells receiving vibroacoustic stimulation were mechanically pushed into denser clusters, producing a distribution of aggregate sizes (measured in mm^2^) that were significantly larger compared to those of controls (*p* = 4.19 *×* 10*^−^*^8^) (Figure 3f). Two days after vibrational treatment, the aggregates in the treated dish had formed organoids that became spontaneously motile (Figure 3g), demonstrating a new automated method for manufacturing motile mucociliary organoids from loose cells.

Chemical delivery was validated at the tissue scale using two concentrations (98µM and 980µM) of the reactive oxygen species H_2_O_2_, delivered with an automated pipetting arm to motile mucociliary organoids (*n* = 3 per concentration), with organoid velocity used as a functional readout (Figure 4a). Exposure to 98µM H_2_O_2_ resulted in increased organoid velocity across the proceeding 20h of observation, while exposure to the higher 980µM concentration of H_2_O_2_ resulted in a decrease in velocity over the same time period (two-way ANOVA, *p* = 0.031, *n* = 3 replicates). These results demonstrate the effect of reactive oxygen species on motile cilia function and support reports in other systems, including mouse airway epithelium [51], illustrating the utility of MOMbot’s chemical delivery hardware in assessing tissue-level outcomes.

Temperature is known to have direct effects on cell, tissue, and organism level-behaviors, from metabolic and developmental rates to heat shock induced protein responses [52]. In the current study, temperature was used to control developmental rate at the organism scale in *Xenopus laevis* embryos across several days of observation. To demonstrate effective thermal coupling between the solid-state heat pump and the liquid filled environment, individual wild-type *Xenopus* embryos at the 2 cell stage were loaded into dishes set at 14°C, 16°C, 18°C, 20°C, and 22°C respectively. Imaging across successive hours allowed tracking of developmental rates and comparisons with established Nieuwkoop and Faber (NF) staging guides (Figure 4b). Embryos in dishes set at 22°C developed about three times faster than siblings raised at 14°C, matching expected values reported in the literature. Temperatures between the upper and lower values demonstrated intermediate developmental rates consistent with established staging predictions. These results confirm that MOMbot’s thermal modulation system can accurately control embryonic development across a wide range of physiologically relevant temperatures for model species.

### 2.4 Autonomous Experimentation

To complement MOMbot’s ability to autonomously execute interventions, it is paired with an artificial intelligence component capable of autonomously designing interventions. This AI component acts by periodically sending interventions it designs to MOMbot, and updates its understanding of the biological target when MOMbot returns the result of that intervention (Figure 5A-i). In this section we investigate whether this AI component has utility: that is, does it send interventions to MOMbot that are more valuable than simply sending interventions drawn randomly from a pre-defined space of possible interventions.

#### Intervention value

The value of an intervention can be defined in many ways. Herein, we determine an intervention to be valuable if slight changes to it cause large changes in the biological target’s response. This follows from the intuition that large contiguous portions of the intervention space may be either too mild, causing no change in behavior, or too severe, causing the same immobilizing effect. The AI component will thus be valuable if it is able to focus interventions in regions of intervention space where the same or similar interventions produce widely varying results (Figure 5A-ii,iii).

#### Intervention space

To estimate whether this AI component is capable, in practice, of converging on valuable interventions for complex interventions and complex biological targets, we restricted ourselves to simple interventions and simple biological targets. Specifically, we restricted ourselves to applying 20 milliamperes of DC current to mucociliary organoids for durations of time between zero and 300 seconds. We chose this restricted intervention space based on our early observations that short applications of 20 milliamperes had no discernable effect on the motility of these organoids, but longer applications permanently impaired their motility.

#### AI component architecture

The AI component itself contains two components. The first component is a search process that seeks valuable interventions to send to MOMbot. The second component is a model that, at any given moment, is more or less certain about how a biological target of interest may respond to a prospective intervention. The search process uses the model’s uncertainty to guide it toward interventions likely to be valuable, if executed by MOMbot.

#### The search process

The search process begins when MOMbot informs it that a new organoid has been loaded into one of its dishes (Figure 5B-i). MOMbot generates a video of the organoid and stores it in the cloud for the search process’s retrieval (Figure 5B-ii). The search process extracts *v*_pre_, the velocity of the organoid’s motion pre-intervention, from the video (Figure 5B-iii). It then feeds *v*_pre_ and the set of possible interventions (Figure 5B-iv) to the model.

#### The model

The model (Figure 5B-v) then outputs *v̂*_post_, a vector that represents its predictions for that organoid’s velocity after any of the possible interventions are applied to it. It also outputs *σ̂*^2^, the estimated variances for each of its predictions. The search process then searches within *σ̂*^2^ to find its maximum value: that is, the model’s maximum uncertainty about how fast the organoid will move, post-intervention (Figure 5B-vi). The candidate intervention which generated this maximal estimated variance is then sent to MOMbot.

#### The intervention result

MOMbot applies the intervention, and returns a video reporting the organoid’s velocity during and after the intervention. This result is used to update the model. The model is updated such that its future predictions about similar interventions resemble the true *v*_post_ and its uncertainty about those predictions are lower. When MOMbot reports that new organoids are available, the cycle repeats.

#### AI component operation

As 29 randomly-chosen electrical interventions had already been applied to 29 organoids during preliminary testing, the AI component was given a warm start: the model was initialized with this data. When MOMbot provided a video of the thirtieth organoid, the search process proposed its first designed intervention. The AI component and MOMbot then alternated designing and intervening upon 55 organoids in eight rounds of parallel experimentation, where the model was updated after every round. This process took ~12 hours over two days (eight 90 minute rounds).

#### Assessing AI component utility

The 29 randomly-chosen interventions were combined with the 55 AIchosen interventions, as well as with 37 interventions chosen manually during preliminary testing, to yield 121 total interventions (Figure 5C-i). The overall response of organoids to these interventions is shown in Figure 5C-ii: for longer duration interventions, post-intervention velocity decreased. To determine whether the 55 AI-generated interventions were converging on valuable interventions—those for which slight changes yielded large changes in organoid behavior—we estimated “ID50” (Immobilization Duration 50), a metric modeled on LD50, a standard toxicological measurement indicating the amount of a substance—administered via food, skin, or inhalation—required to kill 50% of a test population. In ID50 we replace “lethal dose” with “immobilization duration”, yielding the intervention duration below which organoids are generally motile, and above which motility is significantly damaged. If AI-generated interventions are found to approach ID50, then the AI component is finding increasingly valuable interventions. If later AI-generated interventions are no closer to ID50 than earlier ones, the AI component is failing to converge on valuable interventions.

#### ID50 estimation

First, all 121 organoids are binarized according to whether or not they exhibited any detectable motion 40 minutes after their intervention was applied to them. Then, a logistic curve was fit to this binarized post-intervention motility as a function of intervention duration (Figure 5C-iii). ID50, derived from the inflection point in the logistic curve, is estimated to be 148s with a 95% confidence interval of [133, 170]s (Figure 5C-iv). The AI-generated interventions were found to congregate in the model’s region of high uncertainty which is centered on the ID50 (Figure 5C-v). The 55 AI-generated interventions were also found to be statistically significantly closer to the ID50 than the randomly or human-designed interventions are (Figure 5C-vi).

#### AI component conclusions

These results indicate that, after an investigator has pre-determined a space of interventions that MOMbot can execute, and is willing to provide biological targets into MOMbot, the AI component can rapidly converge on interventions that are more valuable than randomly or uniformly-generated ones. Whether this will remain true for more complex biological targets and interventions will be the focus of future studies.

## 3 Discussion

Despite the decades of progress in closed-loop laboratory systems, biology has lacked a platform capable of systematically and autonomously discovering how combined physical stimuli influence the self-organizing dynamics of multiscale living systems. By integrating four distinct intervention modalities, the ability to image biological systems from single cells to vertebrate model organisms, and an active learning algorithm which intelligently selects and executes experiments from an effectively infinite combinatorial space, MOMbot addresses this gap. Here we discuss possible extensions of the system, conceptual drivers of the development of systems of its kind, and broader implications of such a system for synthetic developmental biology.

AI and machine learning are rapidly reshaping biological discovery at the scientific frontier. Beyond their established role in analyzing large-scale imaging and ‘omics datasets, these approaches are poised to drive not only downstream data analysis but also the generation, design, and automated testing of novel experiments across subfields like evolution, development, bioengineering, stem cell biology, aging, and synthetic biology. To advance this vision, the robotic scientist platform described herein is both scale-free and organism-agnostic. The optics, arena configuration, and software architecture described here can be readily extended—without modification—to established model systems including *C. elegans*, *Drosophila*, zebrafish, and axolotl. With the addition of CO_2_ control and off-the-shelf microfluidic components, the system can also be adapted to in vitro platforms, including cultured cell lines, human organoids [53], cancer spheroids [54], and organ-on-chip systems [55]. More broadly, the principles demonstrated here—integrated hardware/software/wetware design, synchronized data pipelines, systematic exploration of experimental variability, and closed-loop hypothesis testing—offer a turnkey framework applicable across diverse areas of biological research.

MOMbot’s software architecture exposes an explicit layer for hardware control, leaving the AI component highly modular and iterable. The active learner was implemented rapidly and consumes few computational resources; more advanced algorithms and machine learning models can be easily swapped in. Additionally, the current age of AI affords new possibilities: LLM-based systems that choose experiments by synthesizing a wide array of data sources (e.g. past experimental data, other relevant datasets, existing literature), can take abstract language-specified research goals as input, or adaptively reconfigure experimental design [56, 57]. It’s currently unclear how these systems will perform, or even by what metrics they should be evaluated. In the future, we intend to iterate on various intelligence architectures on MOMbot.

More conceptually, closing the loop on a laboratory robot precipitates questions about embodied AI and the experimental nature of cognition and intelligence itself. Robot scientists tend to align with the cognitive science frameworks of predictive processing [58] and active inference [59], wherein cognitive agents seek to minimize surprise by taking actions which resolve uncertainty. Closed–loop and high throughput platforms, in addition to advancing scientific understanding, could serve as uniquely suitable testbeds for advancing novel AI methodologies like continual learning, reinforcement learning, the automatic and adaptive construction of world models, and other open questions in embodied artificial intelligence.

As the cost of laboratory hardware falls and the ability and scale of AI grows, we envision a proliferating class of intelligent laboratory robots which continually formulate, test, and revise biological hypotheses in service of human-specified research agendas. Modular platforms like MOMbot can scale experiments across diverse biological substrates in parallel, which is immensely synergistic with computational advances. We view this research and engineering as being positioned at the beginning of a long-term collaboration between human creativity and machine intelligence, whose implications for biology and medicine remain to be seen.

## 4 Methods

### 4.1 Hardware System Architecture

The integrated system takes up a 6ft by 6ft footprint in the lab space. MOMbot’s primary workspace is encased by a protective enclosure, with two doors that remain locked during the system’s usage. The encasing protects lab technicians from the high-velocity gantry system. MOMbot’s workspace contains 5 stations (Section 4.1.1), each with 2 experiment dishes (see Figure S1c), a pipette tip dispenser (Figure S2d), a liquid reservoir (Figure S2c), and a pipette tip discard bin (Figure S2b). Underneath MOMbot’s primary workspace, a chiller unit, a housing unit for electrical wiring, and a housing unit for compute and networking (Figure S8) are all stored (Figure S1a, bottom). On the front of MOMbot is a user interface panel with 4 buttons, a hardware override switch which forces the doors to be open-able and the gantry to shut off, and a toggle switch for resetting the system with a power cycle (Figure S9). The Start button starts the system, the Emergency Stop button cuts power to the system’s hardware, and the Service button unlocks MOMbot’s doors to allow an operator access to the workspace.

#### 4.1.1 Single station

A single MOMbot station is an integrated unit consisting of two independent experimental dishes (Figure S1b, S3). Each station can modulate a single temperature and vibration value for both of its dishes. Otherwise, each dish can independently receive liquid delivery and control electrical stimulus. Each station has a lid containing ambient LED lighting and two cameras, one positioned above each of the two dishes. The station lid contains a Raspberry Pi 5 (RPi5) computer and a custom PCB to network the RPi5, the two cameras, and the ambient LEDs. The bottom of the station contains a voice coil (to provide vibrational stimulus), a thermal plate connected to the system’s chiller, a custom PCB to facilitate the electrical stimulus, and a Raspberry Pi Zero (RPi0) computer to control each of these hardware components. The RPi0 and the RPi5 are connected through a physical UART backchannel, and both are networked to an aggregate network switchboard which connects to the Mombot Core PC (Figure S7).

#### 4.1.2 Imaging

Each station lid on MOMbot consists of two high-resolution (64MP Hawkeye Arducam) cameras which image both dishes on the station (Figure S7). The cameras are embedded in a custom PCB within the station’s lid. The cameras sit 5.2 centimeters above the temperature plate, with an Alstar 0.5x focal reducer sitting 0.33 centimeters in front of the camera. LEDs inside the lid casing illuminate the dishes during imaging through a white acetal sheet diffuser. The Arducam takes 9152x6944 resolution, ~7.4MB images in JPEG format (~80MB raw). Images were compressed at the edge due to network bandwidth constraints.

#### 4.1.3 Chemical delivery system

The chemical delivery system consists of two coupled components: a Zaber LC40 dual motor gantry system and a Hamilton pipetting system (Figure S2). The gantry system can move in the XYZ plane, and the pipette system can move the pipette in the Z axis. Conductive CO-RE pipette tips (Hamilton #235900) sense liquid levels within dishes and chemical reservoirs and dispense volumes of 0.5–10µl as requested by software. When a hardware command requests a chemical stimulus to be delivered to one of the dishes, the wet lab workflow is the following: 1) an indicator light on the system’s light tower indicates a service is needed, and the relevant station’s lid indicates it needs to be removed, 2) an operator opens the system’s doors, removes the lid (imaging is paused during this period), closes the doors, presses start, 3) the gantry acquires a pipette tip at the dispenser, ingests the requested volume of chemical from the reservoir, the chemical is dispensed into the center of the requested dish at a depth of 1mm, the pipette tip is discarded in the discard bin, and the gantry returns to a resting state, and finally 4) the operator opens the doors, replaces the lid, closes the door, and presses start to continue the system’s “normal” operation.

#### 4.1.4 Vibration

Each station delivers vibration stimuli via a linear voice coil motor (Monitcont model number LVCM-044032-02), driven by the bottom PCB’s control software (Figure S4). Four music-wire steel dampener springs (spring constant *k* = 5.6 lbf/in) mechanically isolate vibrations to each station. The stimulus is parameterized by frequency (integer-valued, 7–200 Hz) and amplitude (*A ∈* [0,1]). The voltage applied to the voice coil is *V* = *A · V*_max_(*f*), where *V*_max_ is the piecewise frequency-dependent maximum voltage curve:

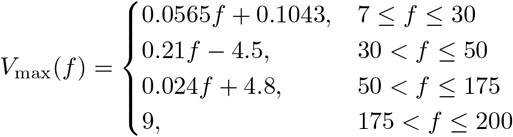

A vibrometer was used to test the worst-case vibration leakage between stations, and stations were found to displace at less than 4% of the vibrating station.

#### 4.1.5 Temperature

A chiller unit (AMS technologies SW10473) is housed beneath MOMbot’s primary workspace (Figure S1a). Tubing is run from the chiller and through each station’s cold plate. One coolant line is used for the entire system, and each station’s temperature is modulated independently by a thermoelectric cooler (TEC), which is controlled by the bottom PCB. The TEC sits between a heat sink (which coolant runs through) and the aluminum plate which petri dishes are loaded onto. The allowable range of temperatures for each station is between 1°C and 30°C with a step size of 1°C. System testing confirmed each station could reach the 1°C setting from the 30°C setting in under 3 minutes.

#### 4.1.6 Electrical

A custom removable electrode ring (Figure S5, right) contains eight iridium-coated titanium electrodes lining the circumference. The electrode ring slots into a pogo pin port on a custom PCB breakout board (Figure S5, left). In the bottom stim apparatus, an RPi0 connects to a more extensive custom PCB which houses the electrical components for modulating each electrode pair independently. The RPi0 houses driver software for the electrode ring. The software can control the direction and the strength of the electric field generated. The field strength ranges between 0 and 20 milliamperes, and the direction of the current ranges between 0 and 360 degrees, at 15 degree increments.

#### 4.1.7 Communication System

MOMbot sits physically in a biology laboratory at Tufts University in Medford, MA, while the AI software can run remotely. Dropbox is used as the communication interface. At max capacity (ten active experiments), MOMbot uploads ~20MB/s of image data to a shared Dropbox folder for real-time computing, while the active learner sends hardware commands (in the form of JSON files) to MOMbot that are executed within seconds. The active learning software constantly monitors the Dropbox for MOMbot to upload image data which are then processed and responded to with hardware commands. Dropbox is an easy-to-use interface where investigators can 1) watch communication between MOMbot and the active learner in real time, 2) use the system without technical knowledge, and 3) have a long-term reliable storage medium and shared workspace for experimental data. Additionally, the Dropbox interface allows the AI to run on a HPC cluster, using file transfer to communicate with MOMbot.

### 4.2 Software System Architecture

MOMbot’s software architecture is split between embedded control software residing on the systems’ onboard computers (the RPi0, RPi5, and the core PC), which manage stimulus execution and image acquisition, and a remote intelligence layer which runs on a remote compute resource and communicates with MOMbot via Dropbox, as described above. The remote layer manages the lifecycle of every dish loaded with biological material as parallel software processes—one per active dish. When a new intervention is required, the process computes pre-intervention velocity from the image stream, queries a variance-driven active learning algorithm to select an intervention maximizing an acquisition function (Section 4.2.2), and writes a structured intervention file that MOMbot’s onboard software executes on hardware. The following sections describe the active learning components in detail. The code repository for MOMbot’s intelligence software is available at https://github.com/kambielawski/mombot_software/.

#### 4.2.1 Interventions

Although MOMbot’s hardware supports multimodal interventions, the autonomous experimentation reported here operated over a single modality—electrical fields at fixed current (20 mA) and fixed E-field angle (270°)—with stimulation duration *d* as the sole free parameter, discretized over 300 candidates spanning 1 to 300 seconds. An intervention is considered valuable if small or no changes in the intervention’s numerical value result in large variation in the behavioral outcome. Again, this is motivated by the intuition that large regions of the intervention search space are “uninteresting” in the sense that they result in the same outcome: either no change in organoid motility (mild interventions) or consistent immobilization (aggressive interventions).

#### 4.2.2 The AI Component

##### Active Learning Search Process

For a given organoid, the active learning algorithm must choose one, of all possible interventions in the search space, to administer to the organoid. The active learning algorithm consists of a forward predictive model and a corresponding variance model that estimates *v*_post_ prediction uncertainty (*σ̂*^2^), which is used as an expected value metric for an acquisition function. The algorithm iteratively hones in on regions of the intervention search space which the predictive model is highly uncertain about, where uncertainty is estimated by the variance model. The following sections describe these active learner components.

##### OLS Forward Prediction

After a pre-observation period of 45 minutes, an organoid’s pre-intervention velocity *v*_pre_ is extracted from MOMbot’s image stream using an image-processing pipeline. A polynomial regression model predicts post-intervention velocity from pre-intervention velocity (*v*_pre_) and a candidate intervention duration (*d*). The raw input vector **x** = [*v*_pre_*, d*] is augmented with nonlinear features to form:

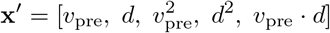

The augmented features are standardized (zero mean, unit variance) using a standard scaler fitted on the training data. We then fit an ordinary least squares (OLS) linear regression on the standardized features to predict *v*_post_:

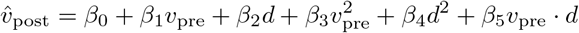

This formulation captures quadratic effects of both input variables and their first-order interaction, while remaining computationally inexpensive to fit as the training set grows incrementally.

##### K-Nearest Neighbor Variance Model

Of the many approaches to active learning, we use a variance-based approach, wherein new interventions are selected based on a variance model [46]. To estimate local outcome uncertainty given a candidate experiment *d* and an initial observation *v*_pre_, we employ a two-stage heteroscedastic variance model [60–62]. First, residuals are computed from the OLS model’s training-set predictions:

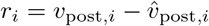

The squared residuals 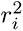 serve as pointwise variance estimates. A *K*-nearest neighbors (KNN) regressor (*K* = 5, distance-weighted) is then fitted to predict 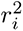 from the raw (non-augmented) input **x** = [*v*_pre_*, d*], separately standardized. For a candidate query point **x***^∗^*, the variance estimate is:

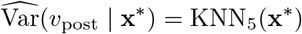

where KNN_5_ returns the distance-weighted average of the five nearest squared residuals. The number of neighbors is clamped to min(*K, n −* 1) where *n* is the current training set size, ensuring the model remains valid in early iterations. Notably, the variance model operates on the raw two-dimensional input space rather than the augmented feature space, so that the KNN distance metric reflects proximity in the physically interpretable (*v*_pre_*, d*) plane (Figure 5C-vi).

##### Acquisition Function

Ultimately, the active learner selects the next experiment by maximizing an upper confidence bound (UCB) acquisition function that balances variance exploitation with density-based exploration [47]. For a given observed *v*_pre_, each candidate duration *d_c_* is scored as follows. First, the predicted variance *σ̂*^2^(*d_c_*) is obtained from the KNN variance model. Second, a density-based exploration bonus is computed:

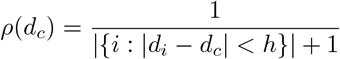

where *h* is a bandwidth parameter (default 50,000 ms) and the count runs over training-set durations *d_i_*. This term is large in regions of the duration axis that are sparsely sampled, providing an additional exploration incentive beyond the variance signal. Both terms are normalized to [0,1] by dividing by their respective maxima across the candidate set, then combined:

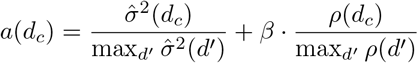

where *β* (= 0.5 in our experiments) controls the exploration–exploitation tradeoff. The selected intervention is *d^∗^* = arg max*_d_ a*(*d_c_*). This acquisition function preferentially samples high-uncertainty regions of the experimental space while also encouraging coverage of underexplored duration ranges, even when the variance model has not yet learned to assign high uncertainty to those regions.

#### 4.2.3 Intervention Execution

Once the active learner selects a duration *d^∗^*, an intervention is constructed specifying a DC electrical stimulation at 20 mA, 0 Hz frequency (DC only), and the selected duration. The intervention is serialized as a JSON command model and transferred to a “pending” directory in Dropbox which is automatically dispatched to the hardware. After the physical execution of the intervention and a post-intervention observation period, the images during this period are uploaded to Dropbox. The resulting *v*_post_ is computed using an automated image processing pipeline consisting of 1) per-frame organoid centroid identification, 2) trajectory extraction, and 3) velocity calculation. *v*_post_ is then appended to the training set for subsequent iterations.

### 4.3 Wetware protocols

#### 4.3.1 Animal Husbandry

All experiments described in the current study were performed under the oversight of the Tufts University Animal Care and Use Committee (IACUC). Procedures involving embryos and living animals were reviewed and approved by the IACUC prior to experimentation, certified under protocols M2020-35 and M2025-40 in compliance with institutional, state, and federal ethical standards for animal welfare. Wild type males and females were sourced to generate all biological material in the present study, including embryos, organoids, and stem cell populations according to standard protocols [63]. Briefly, externally fertilized embryos were de-jellied via a cysteine bath, followed by several washes in 0.1X Marc’s Modified Ringers (MMR, 10x stock: 1M NaCl, 20 mM KCl, 10 mM MgSO4·7H2O, 20 mM CaCl2·2H2O, 50 mM HEPES, pH 7.4). Cohorts of embryos were then raised in a 14 degree incubator, under a 12h/12h light-dark cycle, until they were used for the generation of organoids and stem cells, or placed in the robotic platform for direct experimentation. Embryos were staged according to the *Xenopus* normal table of normal development [64].

#### 4.3.2 Mucociliary organoid generation

Mucociliary organoids were generated from embryonic sources via the standard *Xenopus* animal cap assay [65]. Cohorts of Nieuwkoop and Faber stage 10 *Xenopus* embryos had their vitelline membranes removed manually with a pair of microsurgery forceps, before being transferred to a petri dish containing 0.75x MMR and 1µg/ml gentamicin, lined with a 1% agarose substrate. The animal cap, composed of the topmost pigmented portion of the developing embryo, was then microdissected using surgical forceps, and inverted on the agar substrate, where it was allowed to heal into a spheroid over the course of 1h at room temperature. The remaining portions of operated embryos were then discarded, while the spheroids were transferred to the robotic platform or stored in a 14°C incubator until experimentation. Developing organoids begin as a mass of undifferentiated stem cell tissue, which over the course of three days then self-organizes into a functional mucociliary organoid with motile cilia lining the external surface of the tissue. Development does not require additional chemical/genetic factors or cell culture media, as the tissue is able to consume maternally loaded yolk-based energy sources over its 10-14 day lifespan. All experiments used single organoids in each station, with media being refreshed daily as needed.

#### 4.3.3 Epidermal stem cell generation

All epidermal stem cell populations were harvested from developing *Xenopus* embryos at Nieuwkoop and Faber stage 10. The vitelline membrane of individual embryos were removed with a pair of microsurgery forceps, before being transferred to a fresh 1% agarose-coated Petri dish containing calciumand magnesiumfree dissociation medium (50.3 mM NaCl, 0.7 mM KCl, 9.2 mM Na_2_HPO_4_, 0.9 mM KH_2_PO_4_, 2.4 mM NaHCO_3_, 1.0 mM edetic acid, pH 7.3) [66]. The animal cap, as described in the section above, was then manually removed and left undisturbed in the dissociation medium for 5 minutes. In this solution, the pigmented outer ectoderm layer remains intact. This layer was carefully separated from the underlying stem cells using surgical forceps and discarded. The remaining tissue was then mechanically agitated by pipetting until it was fully dissociated into single cells. Cells from 30 embryos were pooled to create a combined cell suspension. The pooled cells were collected and transferred into a sterile Eppendorf tube containing 100µl of 0.75× MMR, then gently mixed by pipetting up and down five additional times to produce a uniform stem cell suspension. Using a clean 200µl pipette, the suspension was placed into a station within the robotic platform containing a 1% agarose-coated Petri dish and 10ml 0.75× MMR. The angle and speed of dispensing determined the final cell density in the dish. After plating, cells were allowed to settle for 1 minute before initiating experiments.

#### 4.3.4 MOMbot wetware workflow

Biological experiments using the MOMbot platform began by placing 60mm polystyrene Petri dishes into each station, lined with 3ml of 1.0% agarose made with 0.1X or 0.75X MMR depending on whether experiments used embryos or organoids/cells respectively. E-stim inserts were then placed into each Petri dish, which contacted the mainboard via pogo pin connectors and were locked into place mechanically. Agar levels ensured contact between the base of the e-stim insert and the substrate, ensuring even current levels across the surface of the arena. 10ml of 0.1X or 0.75X MMR (for embryos or organoids/cells respectively), pH 7.4 was then added to each station via serological pipette. Biological tissue in the form of single embryos, organoids, or several thousand loose cells, were then deposited into the center of each arena via transfer pipette. Once loaded, the optics were seated on top of each station, and the experimental temperature was set for each experiment. For experiments running longer than 12h, liquid levels were manually maintained by adding fresh media 2x daily, between image capture events.

Upon completion of an experiment, remaining biological materials were collected, frozen, and stored for institutional biological waste disposal. Media was discarded, except in cases where chem-stim delivered hazardous chemicals, in which case all remaining media was collected for weekly waste disposal. Agar coated petri dishes were disposed of in standard mixed waste receptacles. E-stim inserts were cleaned between each experiment; first by a wash in 70% ethanol, followed by a wash in distilled water before drying via blotting with Kimwipes. All stations were inspected between experiments to verify no water spilling or ingress outside of the e-stim ring and Petri dish. Any overflow was removed with absorbent towels followed by a wash with distilled water to remove any salt residue.

## Acknowledgements

This research was supported by the Engineer Research and Development Center – Cold Regions Research and Engineering Laboratory (ERDCCRREL) under Contract No. W913E524C0012. The content of the information does not necessarily reflect the position or policy of the government, and no official endorsement should be inferred. We would also like to thank Martin Schwalm and Thomas Varley for their input on experimental design.

## Author Contributions

Conceptualization: D.B., J.B, K.B., and M.L.; Methodology: D.B., J.B, K.B., S.B., N.G, and K.S.; Experimental investigation: D.B. and K.B.; Data analysis: D.B., K.B., S.B, N.G., and K.S; Data interpretation: D.B., J.B, K.B., S.B., N.G, M.L., J.L., and K.S.; Resources and valuable inputs: N.G. and J.L.; Original draft: D.B. and K.B. All authors reviewed, edited and approved the manuscript.

## Code Availability

https://github.com/kambielawski/mombot_software/

## Supplementary Information

**Figure S1:**
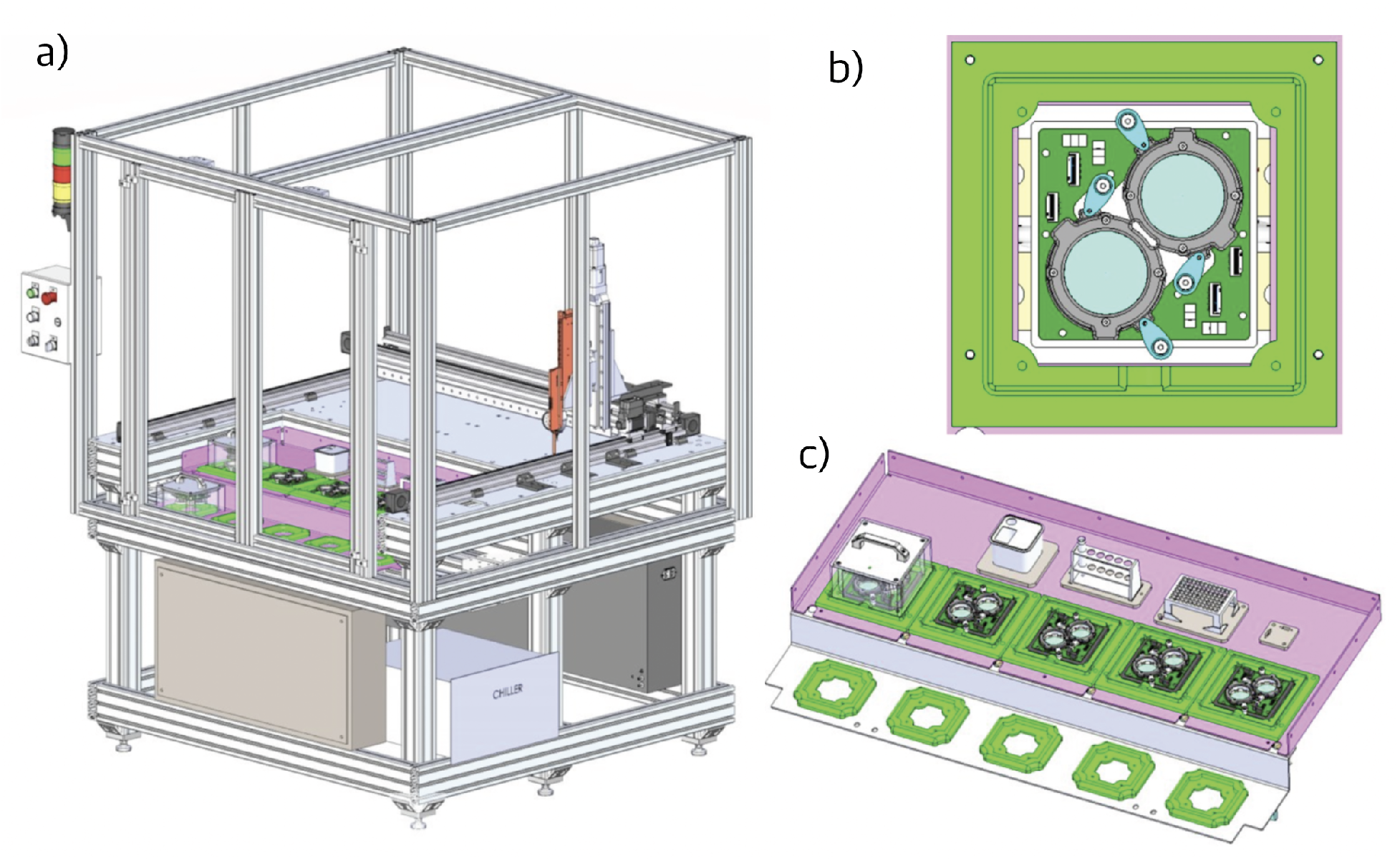
System overview CAD schematics. a) Schematic for MOMbot system. 6ft x 6ft laboratory layout. Primary workspace (see c) is encased for safe operation of the semi-automated pipette system (in orange; see Figure S2). b) Top-down view of one station, with two experimental dishes. c) MOMbot primary workspace, consisting of 5 stations. Above the stations, from left to right: pipette tip discard bin, chemical reservoirs, pipette tip dispenser.

**Figure S2:**
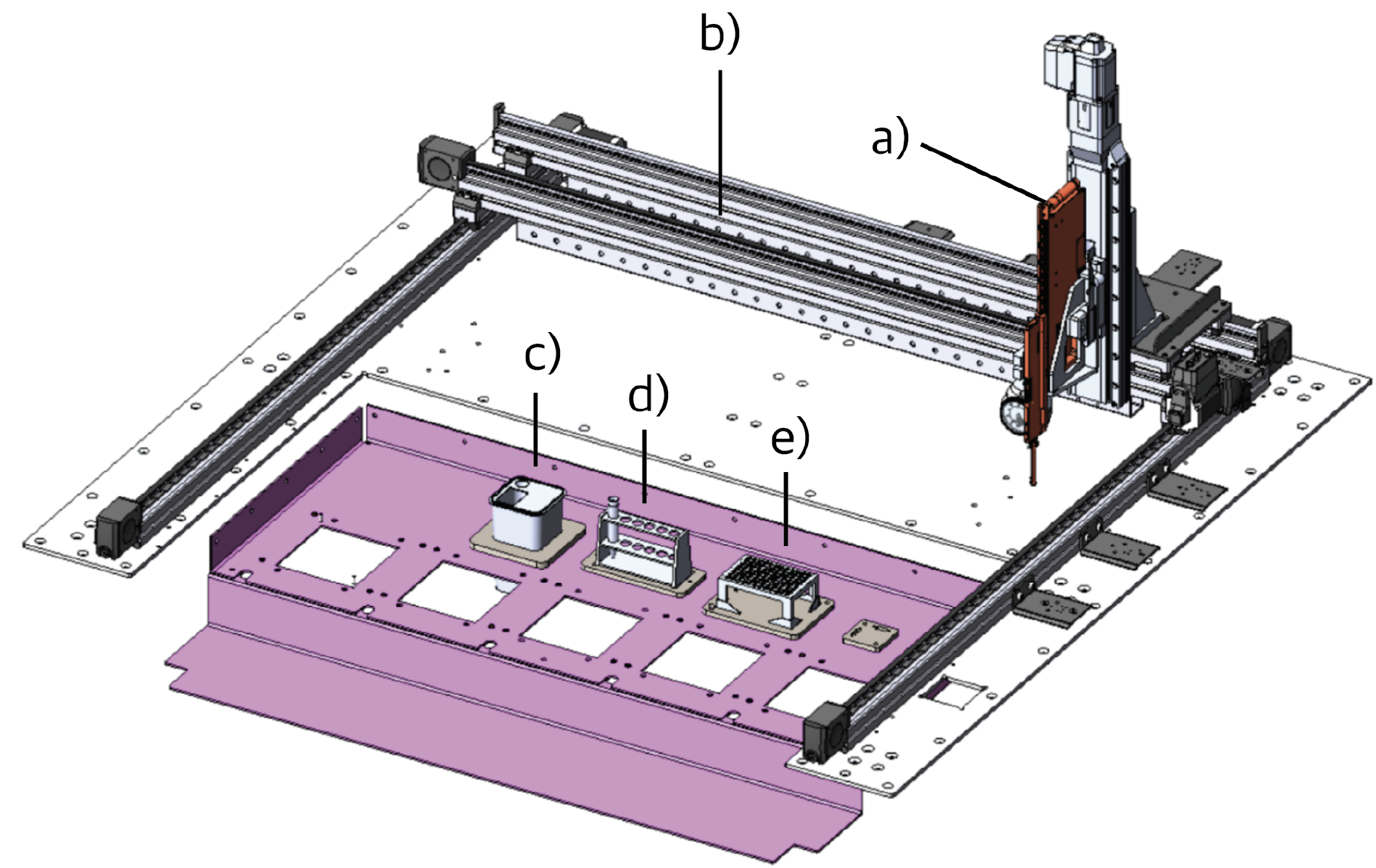
Gantry/pipette subsystem. a) Hamilton pipetting system, capable of moving in the Z axis. b) Linear gantry rails, capable of moving in the X-Y axes. c) Pipette tip discard bin. d) Chemical reservoirs.e) Pipette tip dispenser.

**Figure S3:**
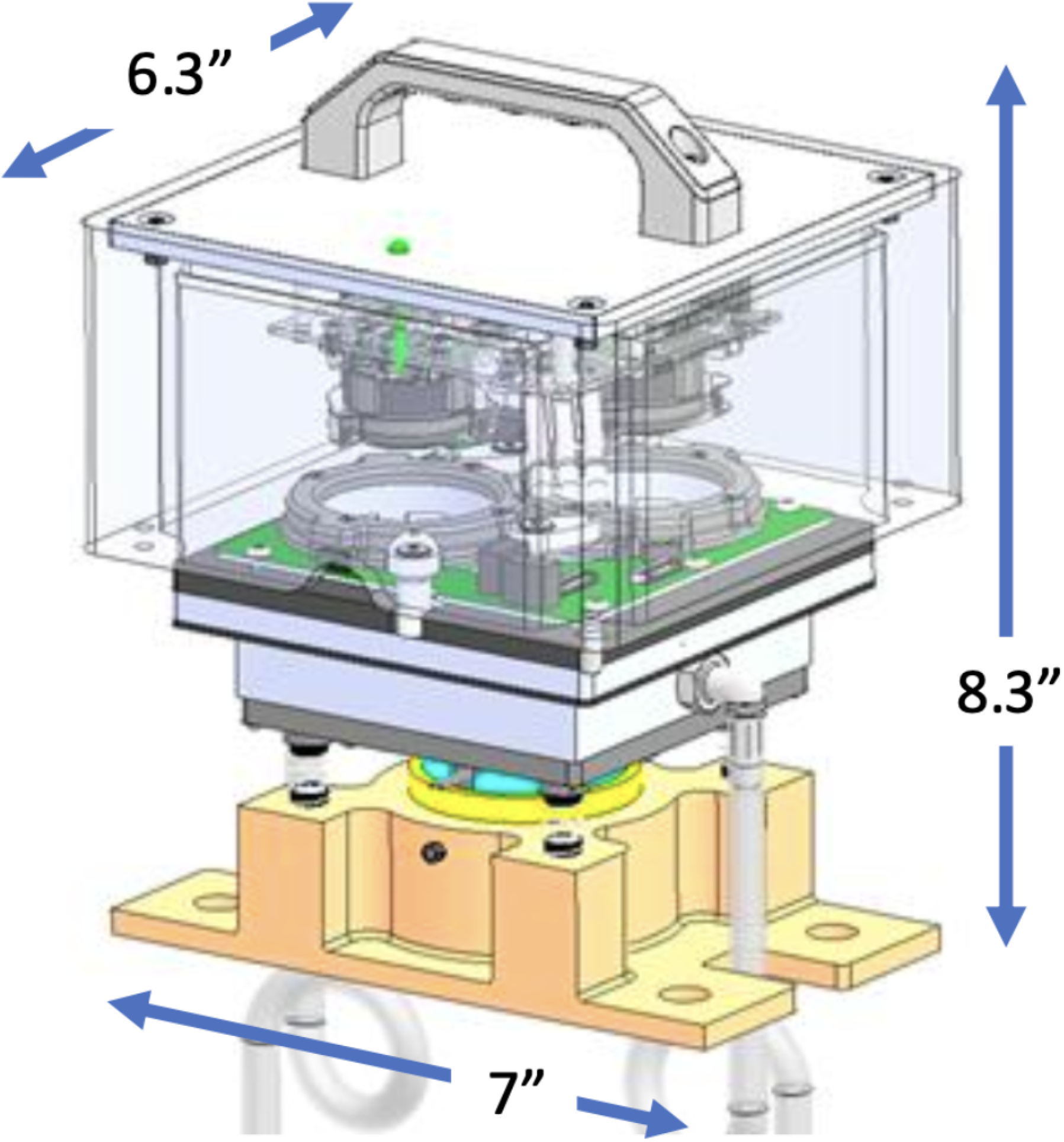
Station subsystem w/ dimensions. Five of these subsystems operate independently and in parallel in MOMbot’s primary workspace.

**Figure S4:**
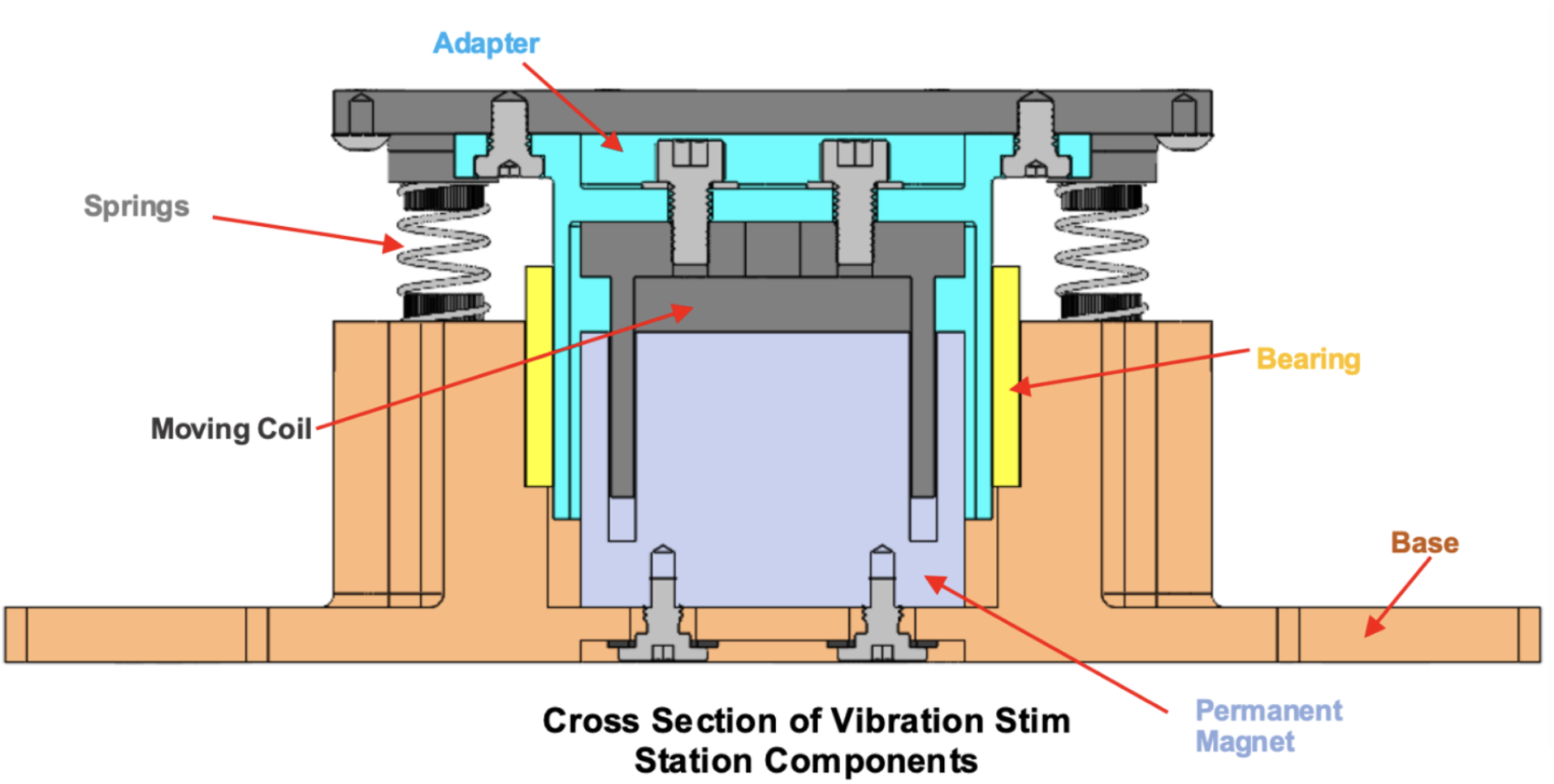
Annotated vibration subsystem.

**Figure S5:**
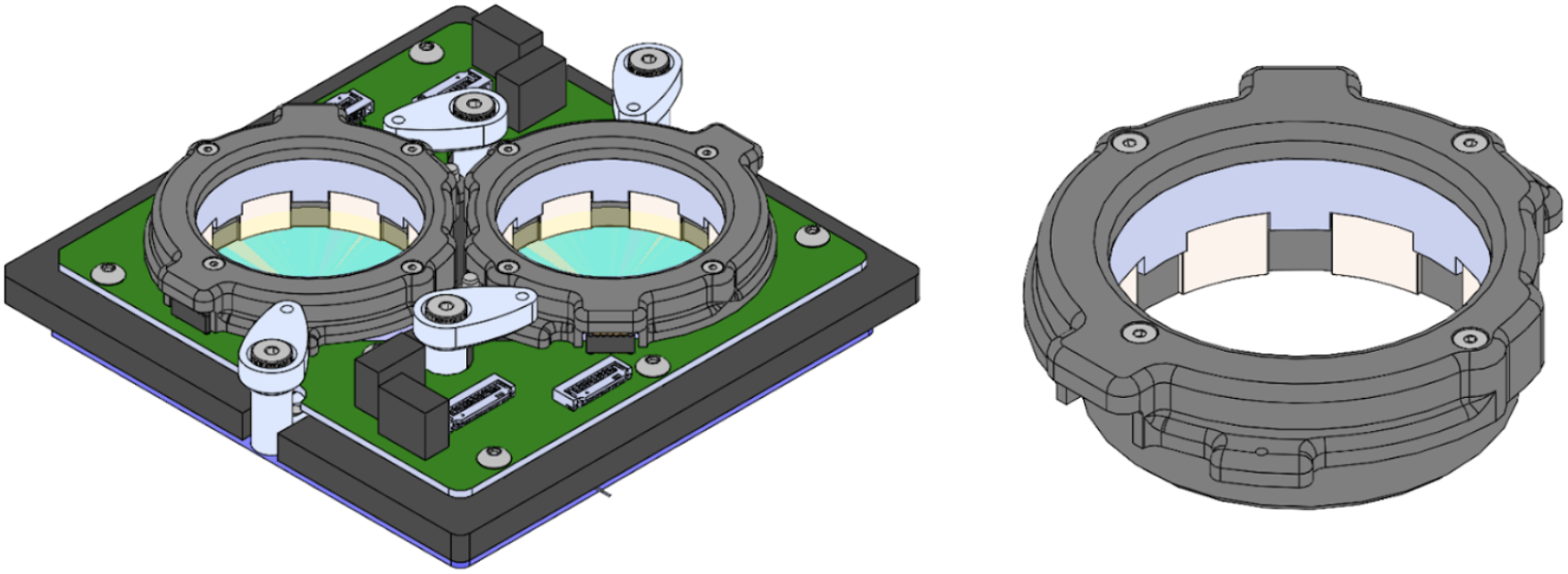
Electrical stim subsystem. Left: CAD schematic of one station, with electrode ring inserts inserted. Right: Electrode ring insert.

**Figure S6:**
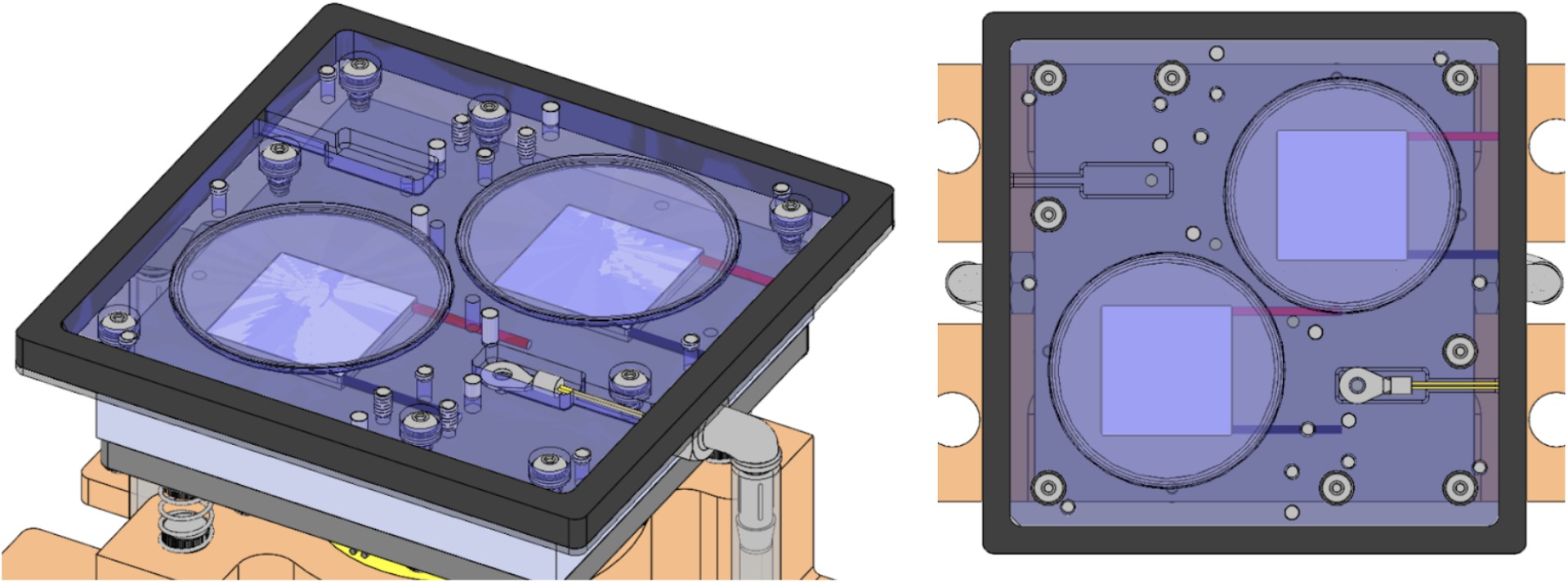
Thermal subsystem. Squares under each dish are thermo-electric coolers (TECs). Yellow wire is a thermistor. The station’s shared aluminum plate (which experiment dishes sit on) is transparent blue.

**Figure S7:**
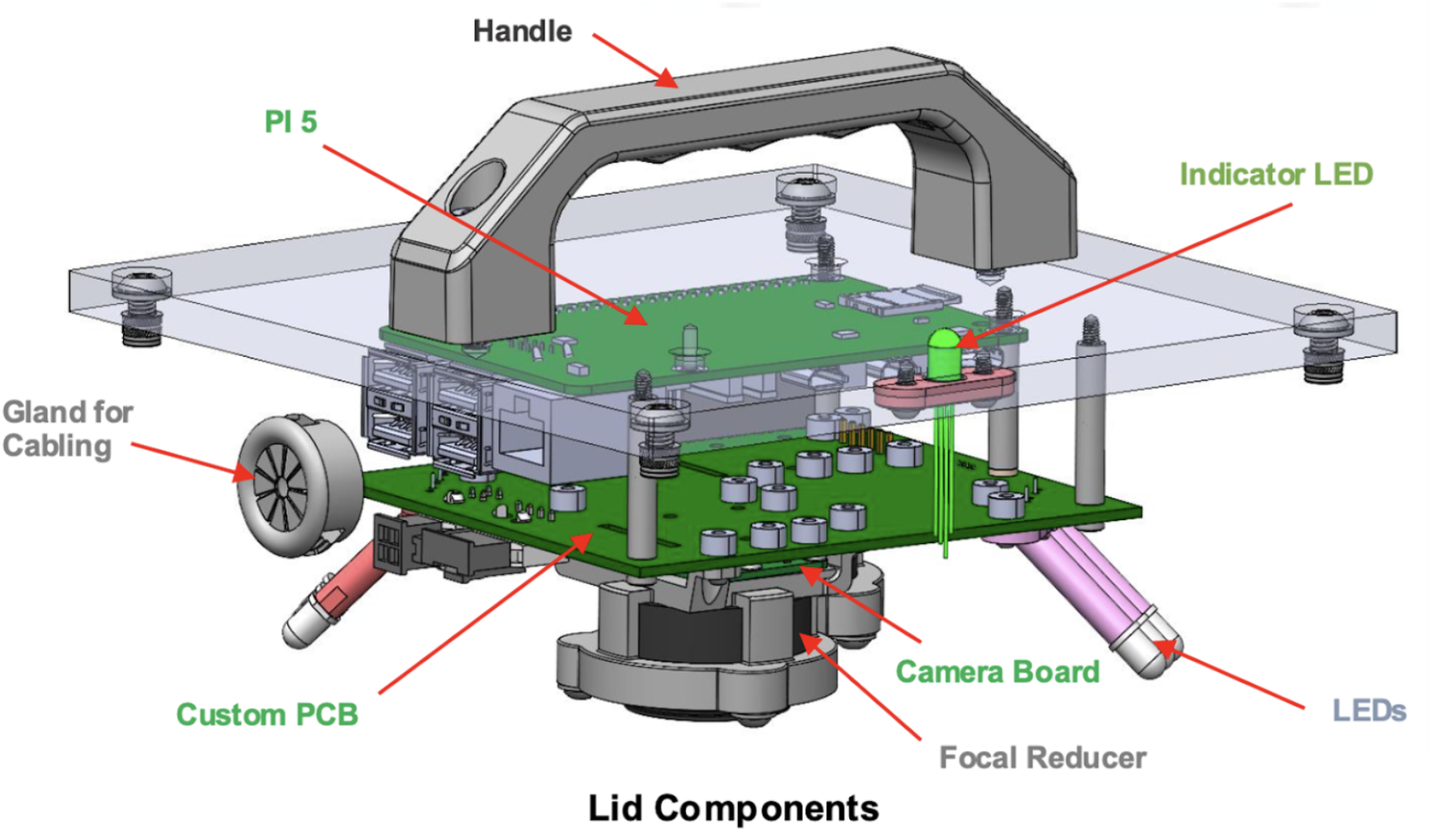
Annotated lid subsystem.

**Figure S8:**
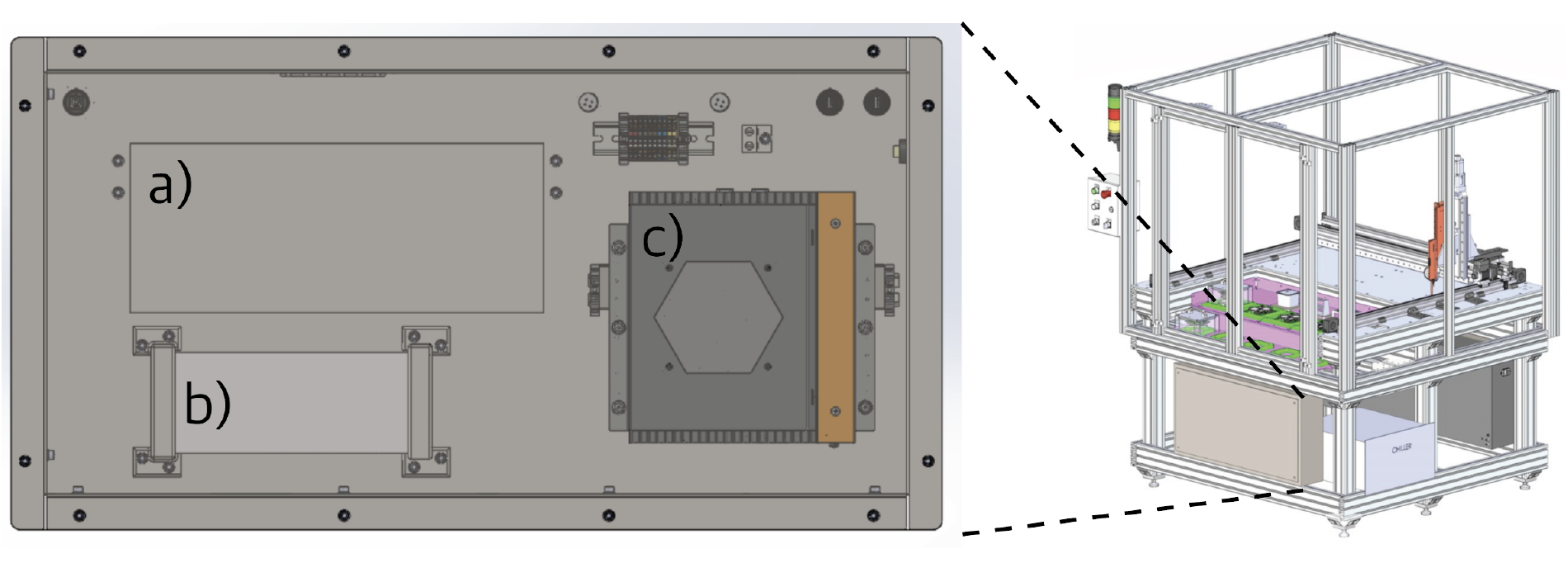
Front Panel. a) Network switchboard integrating signals from all 5 stations. b) Liquid handling controller. c) Core MOMbot PC: Karbon 801 by OnLogic.

**Figure S9:**
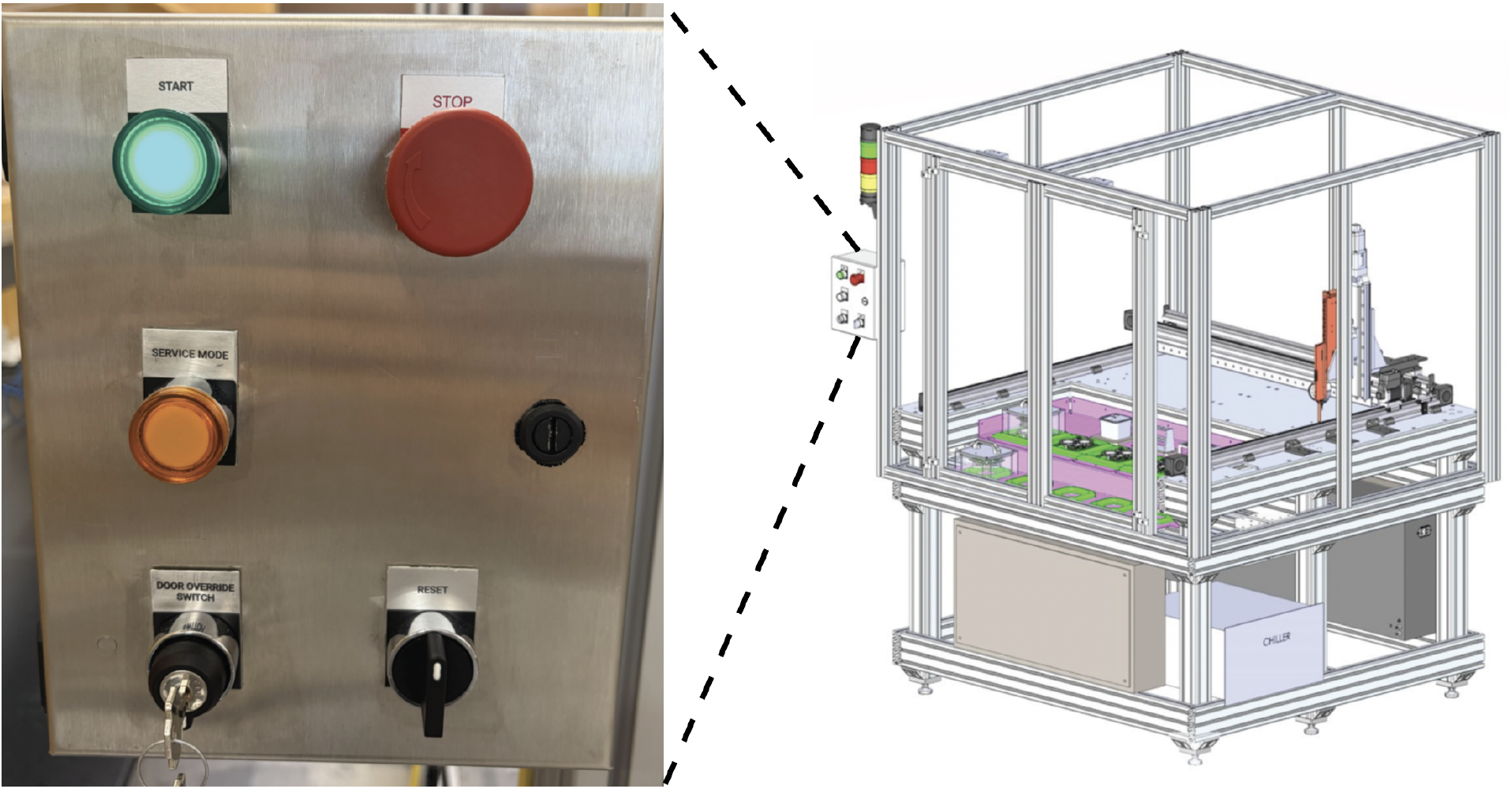
MOMbot operator panel. The green “Start” button (top-left) puts MOMbot into a default state, with doors locked and gantry system active. The Emergency Stop button (top-right) kills power to the system in case of emergency. The “Service” button (middle-left) puts MOMbot into a service mode, wherein MOMbot’s doors unlock and the gantry system is disabled, for safe operator access to the stations in the primary workspace. The Door override switch (bottom-left) can allow the operator access to the inner workspace; this switch also kills power to the gantry system for safe access. The reset toggle (bottom-right) is triggered to reboot the system if it is powered off.

## Notes

https://github.com/kambielawski/mombot_software/

### Competing Interest Statement

The authors have declared no competing interest.

## References

1. Abramson, J. et al. Accurate Structure Prediction of Biomolecular Interactions with AlphaFold 3. Nature 630, 493–500. issn: 1476-4687. https://www.nature.com/articles/s41586-024-07487-w (2026) (June 2024).

2. King, R. D. et al. Functional genomic hypothesis generation and experimentation by a robot scientist. eng. Nature 427, 247–252. issn: 1476-4687 (Jan. 2004).

3. Saitoh, S. & Yoshimori, T. Fully Automated Laboratory Robotic System for Automating Sample Preparation and Analysis to Reduce Cost and Time in Drug Development Process. JALA: Journal of the Association for Laboratory Automation 13, 265–274. issn: 1535-5535. https://journals.sagepub.com/doi/10.1016/j.jala.2008.07.001 (2025) (Oct. 2008).

4. Coley, C. W., Eyke, N. S. & Jensen, K. F. Autonomous Discovery in the Chemical Sciences Part I: Progress. en. Angewandte Chemie International Edition 59, 22858–22893. issn: 1433-7851, 1521-3773. https://onlinelibrary.wiley.com/doi/10.1002/anie.201909987 2024) (Dec. 2020).

5. Burger, B., et al. A mobile robotic chemist. en. Nature 583. Publisher: Nature Publishing Group, 237–241. issn: 1476-4687. https://www.nature.com/articles/s41586-020-2442-2 (2025) (July 2020).

6. Volk, A. A. et al. AlphaFlow: autonomous discovery and optimization of multi-step chemistry using a self-driven fluidic lab guided by reinforcement learning. en. Nature Communications 14, 1403. issn: 2041-1723. https://www.nature.com/articles/s41467-023-37139-y (2024) (Mar. 2023).

7. Wigley, P. B., et al. Fast machine-learning online optimization of ultra-cold-atom experiments. en. Scientific Reports 6, 25890. issn: 2045-2322. https://www.nature.com/articles/srep25890 (2025) (May 2016).

8. Roccapriore, K. M., Dyck, O., Oxley, M. P., Ziatdinov, M. & Kalinin, S. V. Automated Experiment in 4D-STEM: Exploring Emergent Physics and Structural Behaviors. eng. ACS nano 16, 7605–7614. issn: 1936-086X (May 2022).

9. Xue, D., et al. Accelerated search for materials with targeted properties by adaptive design. en. Nature Communications 7. Publisher: Nature Publishing Group, 11241. issn: 2041-1723. https://www.nature.com/articles/ncomms11241 (2025) (Apr. 2016).

10. Nikolaev, P. et al. Autonomy in materials research: a case study in carbon nanotube growth. en. npj Computational Materials **2**. Publisher: Nature Publishing Group, 1–6. issn: 2057-3960. https://www.nature.com/articles/npjcompumats201631 (2025) (Oct. 2016).

11. Tabor, D. P. et al. Accelerating the discovery of materials for clean energy in the era of smart automation. en. Nature Reviews Materials 3, 5–20. issn: 2058-8437. https://www.nature.com/articles/s41578-018-0005-z (2025) (Apr. 2018).

12. Szymanski, N. J. et al. An autonomous laboratory for the accelerated synthesis of novel materials. en. Nature 624, 86–91. issn: 0028-0836, 1476-4687. https://www.nature.com/articles/s41586-023-06734-w (2024) (Dec. 2023).

13. Abolhasani, M. & Kumacheva, E. The rise of self-driving labs in chemical and materials sciences. en. Nature Synthesis 2, 483–492. issn: 2731-0582. https://www.nature.com/articles/s44160-022-00231-0 (2025) (Jan. 2023).

14. Hu, R. et al. Protein Engineering via Bayesian Optimization-Guided Evolutionary Algorithm and Robotic Experiments. Briefings in Bioinformatics 24, bbac570. issn: 1477-4054. PMID: 36562723 (Jan. 19, 2023).

15. Yang, J. et al. Active Learning-Assisted Directed Evolution. Nature Communications 16, 714. issn: 2041-1723. https://www.nature.com/articles/s41467-025-55987-8 (2025) (Jan. 16, 2025).

16. Langley, P. Integrated Systems for Computational Scientific Discovery. Proceedings of the AAAI Conference on Artificial Intelligence 38, 22598–22606. issn: 2374-3468, 2159-5399. https://ojs.aaai.org/index.php/AAAI/article/view/30269 (2024) (Mar. 24, 2024).

17. Gower, A. H., et al. The Use of AI-Robotic Systems for Scientific Discovery version 1. arXiv: 2406.17835 [cs]. http://arxiv.org/abs/2406.17835 (2024). Pre-published.

18. Cooper, A. I. et al. Accelerating Discovery in Natural Science Laboratories with AI and Robotics: Perspectives and Challenges from the 2024 IEEE ICRA Workshop, Yokohama, Japan version 1. arXiv: 2501.06847 [cs]. http://arxiv.org/abs/2501.06847 (2026). Pre-published.

19. Zournas, A. et al. Machine learning-led semi-automated medium optimization reveals salt as key for flaviolin production in Pseudomonas putida. en. Communications Biology 8. Publisher: Nature Publishing Group, 630. issn: 2399-3642. https://www.nature.com/articles/s42003025-080392 (2025) (Apr. 2025).

20. Kanda, G. N., et al. Robotic search for optimal cell culture in regenerative medicine. en. eLife 11, e77007. issn: 2050-084X. https://elifesciences.org/articles/77007 (2025) (June 2022).

21. Blackiston, D., Shomrat, T., Nicolas, C. L., Granata, C. & Levin, M. A Second-Generation Device for Automated Training and Quantitative Behavior Analyses of Molecularly-Tractable Model Organisms. en. PLOS ONE 5. Publisher: Public Library of Science, e14370. issn: 1932-6203. https://journals.plos.org/plosone/article?id=10.1371/journal.pone.0014370 (2025) (Dec. 2010).

22. Savall, J., Ho, E. T. W., Huang, C., Maxey, J. R. & Schnitzer, M. J. Dexterous robotic manipulation of alert adult Drosophila for high-content experimentation. en. Nature Methods 12, 657–660. issn: 1548-7091, 1548-7105. https://www.nature.com/articles/nmeth.3410 (2025) (July 2015).

23. Lin, Y., Silverman-Dultz, A., Bailey, M. & Cohen, D. J. A programmable, open-source robot that scratches cultured tissues to investigate cell migration, healing, and tissue sculpting. en. Cell Reports Methods 4, 100915. issn: 26672375. https://linkinghub.elsevier.com/retrieve/pii/S2667237524003059 (2025) (Dec. 2024).

24. Tamasi, M. J. et al. Machine Learning on a Robotic Platform for the Design of Polymer-Protein Hybrids. eng. Advanced Materials (Deerfield Beach, Fla.) 34, e2201809. issn: 1521-4095 (July 2022).

25. Dubey, A. & Saint-Jeannet, J.-P. Modeling Human Craniofacial Disorders in Xenopus. Current Pathobiology Reports 5, 79–92. issn: 2167-485X. PMID: 28255527 (Mar. 2017).

26. Hardwick, L. J. A. & Philpott, A. Xenopus Models of Cancer: Expanding the Oncologist’s Toolbox. Frontiers in Physiology 9. issn: 1664-042X. https://www.frontiersin.org/journals/physiology/articles/10.3389/fphys.2018.01660/full (2026) (Nov. 27, 2018).

27. Exner, C. R. T. & Willsey, H. R. Xenopus Leads the Way: Frogs as a Pioneering Model to Understand the Human Brain. Genesis (New York, N.Y. : 2000) 59, e23405. issn: 1526-954X. PMID: 33369095. https://pmc.ncbi.nlm.nih.gov/articles/PMC8130472/ (2026) (Feb. 2021).

28. Walentek, P. Xenopus Epidermal and Endodermal Epithelia as Models for Mucociliary Epithelial Evolution, Disease, and Metaplasia. Genesis 59, e23406. issn: 1526-968X. PMID: 33400364 (Feb. 2021).

29. Walentek, P. & Quigley, I. K. What We Can Learn from a Tadpole about Ciliopathies and Airway Diseases – Using Systems Biology in Xenopus to Study Cilia and Mucociliary Epithelia. Genesis (New York, N.Y. : 2000) 55, 10.1002/dvg.23001. issn: 1526-954X. PMID: 28095645. https://pmc.ncbi.nlm.nih.gov/articles/PMC5276738/ (2026) (Jan. 2017).

30. Dubaissi, E. & Papalopulu, N. Embryonic Frog Epidermis: A Model for the Study of Cell-Cell Interactions in the Development of Mucociliary Disease. Disease Models & Mechanisms 4, 179–192. issn: 1754-8411. PMID: 21183475 (Mar. 2011).

31. Blackiston, D. et al. A Cellular Platform for the Development of Synthetic Living Machines. Science Robotics 6, eabf1571. https://www.science.org/doi/10.1126/scirobotics.abf1571 (2026) (Mar. 31, 2021).

32. Kriegman, S., Blackiston, D., Levin, M. & Bongard, J. Kinematic self-replication in reconfigurable organisms. Proceedings of the National Academy of Sciences 118. Publisher: Proceedings of the National Academy of Sciences, e2112672118. https://www.pnas.org/doi/10.1073/pnas.2112672118 (2025) (Dec. 2021).

33. Li, H. et al. Material-Engineered Bioartificial Microorganisms Enabling Efficient Scavenging of Waterborne Viruses. Nature Communications 14, 4658. issn: 2041-1723. https://www.nature.com/articles/s41467-023-40397-5 (2026) (Aug. 3, 2023).

34. Raman, R. Biofabrication of Living Actuators. Annual Review of Biomedical Engineering 26, 223–245. issn: 1523-9829, 1545-4274. https://www.annualreviews.org/content/journals/10.1146/annurev-bioeng-110122-013805 (2026) (Volume 26, 2024 July 3, 2024).

35. Ricotti, L. et al. Biohybrid Actuators for Robotics: A Review of Devices Actuated by Living Cells. Science Robotics 2, eaaq0495. https://www.science.org/doi/10.1126/scirobotics.aaq0495 (2026) (Nov. 29, 2017).

36. Yap, T. F., Liu, Z., Rajappan, A., Shimokusu, T. J. & Preston, D. J. Necrobotics: Biotic Materials as Ready-to-Use Actuators. Advanced Science 9, e2201174. issn: 2198-3844. PMID: 35875913 (Oct. 2022).

37. Kriegman, S., Blackiston, D., Levin, M. & Bongard, J. A scalable pipeline for designing reconfigurable organisms. Proceedings of the National Academy of Sciences 117, 1853–1859. issn: 0027-8424. eprint: https://www.pnas.org/content/117/4/1853.full.pdf. https://www.pnas.org/content/117/4/185 (2020).

38. Bhattaram, D. et al. AggreBots: Configuring CiliaBots through Guided, Modular Tissue Aggregation. Science Advances 11, eadx4176. https://www.science.org/doi/10.1126/sciadv.adx4176 (2026) (Sept. 26, 2025).

39. Chen, C. et al. 3D-printed Microfluidic Devices: Fabrication, Advantages and Limitations—a Mini Review. Analytical methods : advancing methods and applications 8, 6005–6012. issn: 1759-9660. PMID: 27617038. https://pmc.ncbi.nlm.nih.gov/articles/PMC5012532/ (2026) (Aug. 21, 2016).

40. Grebenyuk, S. et al. Large-scale perfused tissues via synthetic 3D soft microfluidics. Nature Communications 14, 1–18. https://ideas.repec.org/a/nat/natcom/v14y2023i1d10.1038_s41467-022-35619-1.html (Dec. 2023).

41. Bai, J. et al. From Platform to Knowledge Graph: Evolution of Laboratory Automation. JACS Au 2, 292–309. 10.1021/jacsau.1c00438 (2026) (Feb. 28, 2022).

42. Antonios, K., Croxatto, A. & Culbreath, K. Current State of Laboratory Automation in Clinical Microbiology Laboratory. Clinical Chemistry 68, 99–114. issn: 1530-8561. PMID: 34969105 (Dec. 30, 2021).

43. Darvish, K., et al. ORGANA: A Robotic Assistant for Automated Chemistry Experimentation and Characterization arXiv: 2401.06949 [cs]. http://arxiv.org/abs/2401.06949 (2026). Pre-published.

44. Seung, H. S., Opper, M. & Sompolinsky, H. Query by committee en. in Proceedings of the fifth annual workshop on Computational learning theory (ACM, Pittsburgh Pennsylvania USA, July 1992), 287–294. isbn: 978-0-89791-497-0. https://dl.acm.org/doi/10.1145/130385.130417 (2025).

45. Jones, D. R. & Schonlau, M. Efficient Global Optimization of Expensive Black-Box Functions. en.

46. Settles, B. Active Learning Literature Survey. en.

47. Shahriari, B., Swersky, K., Wang, Z., Adams, R. P. & de Freitas, N. Taking the Human Out of the Loop: A Review of Bayesian Optimization. Proceedings of the IEEE 104. Conference Name: Proceedings of the IEEE, 148–175. issn: 1558-2256. https://ieeexplore.ieee.org/document/7352306/?arnumber=7352306 (2025) (Jan. 2016).

48. Freedman, L. P., Cockburn, I. M. & Simcoe, T. S. The Economics of Reproducibility in Preclinical Research. PLOS Biology 13, e1002165. issn: 1545-7885. https://journals.plos.org/plosbiology/=article?id=10.1371/journal.pbio.1002165 (2026) (June 9, 2015).

49. Kim, D. & Kwon, S. Vibrational Stress Affects Extracellular Signal-Regulated Kinases Activation and Cytoskeleton Structure in Human Keratinocytes. PLOS ONE 15, e0231174. issn: 1932-6203. https://journals.plos.org/plosone/article? id=10.1371/journal.pone.0231174 (2026) (Apr. 8, 2020).

50. Guo, F. et al. Controlling Cell-Cell Interactions Using Surface Acoustic Waves. Proceedings of the National Academy of Sciences of the United States of America 112, 43–48. issn: 1091-6490. PMID: 25535339 (Jan. 6, 2015).

51. Francis, R. The Effects of Acute Hydrogen Peroxide Exposure on Respiratory Cilia Motility and Viability. PeerJ 11, e14899 (Feb. 27, 2023).

52. Knapp, B. D. & Huang, K. C. The Effects of Temperature on Cellular Physiology. Annual Review of Biophysics 51, 499–526. issn: 1936-122X, 1936-1238. https://www.annualreviews.org/doi/10.1146/annurev-biophys-112221-074832 (2026) (May 9, 2022).

53. Kim, J., Koo, B.-K. & Knoblich, J. A. Human Organoids: Model Systems for Human Biology and Medicine. Nature Reviews Molecular Cell Biology 21, 571–584. issn: 1471-0080. https://www.nature.com/articles/s41580-020-0259-3 (2026) (Oct. 2020).

54. Ishiguro, T., et al. Tumor-Derived Spheroids: Relevance to Cancer Stem Cells and Clinical Applications. Cancer Science 108, 283–289. issn: 1349-7006. PMID: 28064442 (Mar. 2017).

55. Ingber, D. E. Human Organs-on-Chips for Disease Modelling, Drug Development and Personalized Medicine. Nature Reviews Genetics 23, 467–491. issn: 1471-0064. https://www.nature.com/articles/s41576-022-00466-9 (2026) (Aug. 2022).

56. Smith, A. A. et al. Using a GPT-5-driven Autonomous Lab to Optimize the Cost and Titer of Cell-Free Protein Synthesis https://www.biorxiv.org/content/10.64898/2026.02.05.703998v1 (2026). Pre-published.

57. Swanson, K., Wu, W., Bulaong, N. L., Pak, J. E. & Zou, J. The Virtual Lab of AI Agents Designs New SARS-CoV-2 Nanobodies. Nature 646, 716–723. issn: 1476-4687. https://www.nature.com/articles/s41586-025-09442-9 (2026) (Oct. 2025).

58. Ciria, A., Schillaci, G., Pezzulo, G., Hafner, V. V. & Lara, B. Predictive Processing in Cognitive Robotics: A Review arXiv: 2101.06611 [cs]. http://arxiv.org/abs/2101.06611 (2026). Pre-published.

59. Friston, K. et al. Active Inference and Learning. Neuroscience and Biobehavioral Reviews 68, 862–879. issn: 0149-7634 (Sept. 2016).

60. Davidian, M. & Carroll, R. J. Variance Function Estimation. Journal of the American Statistical Association 82, 1079–1091. issn: 0162-1459. https://www.tandfonline.com/doi/abs/10.1080/01621459.1987.10478543 (2026) (Dec. 1, 1987).

61. Muller, H.-G. & Stadtmuller, U. Estimation of Heteroscedasticity in Regression Analysis. The Annals of Statistics 15, 610–625. issn: 0090-5364, 2168-8966. https://projecteuclid.org/journals/annalsofstatistics/volume-15/issue-2/EstimationofHeteroscedasticityinRegressionAnalysis/10.1214/aos/1176350364.full (2026) (June 1987).

62. Liitïainen, E., Corona, F. & Lendasse, A. Residual Variance Estimation Using a Nearest Neighbor Statistic. Journal of Multivariate Analysis 101, 811–823. issn: 0047-259X. https://www.sciencedirect.com/science/article/pii/S0047259X10000035 (2026) (Apr. 1, 2010).

63. Sive, H. L., Grainger, R. M. & Harland, R. M. Early development of Xenopus laevis (Cold Spring Harbor Laboratory Press, New York, NY, Mar. 2010).

64. Normal table of Xenopus laevis (daudin) (eds Faber, J. & Nieuwkoop, P. D.) (CRC Press, Boca Raton, FL, June 1994).

65. Green, J. The Animal Cap Assay. Methods in molecular biology (Clifton, N.J.) 127, 1–13. issn: 159259-678-9 (Feb. 1, 1999).

66. Ariizumi, T. et al. Isolation and Differentiation of Xenopus Animal Cap Cells. Current Protocols in Stem Cell Biology **Chapter** 1, Unit 1D.5. issn: 1938-8969. PMID: 19382122 (Apr. 2009).

